# Genotyping Polyploids from Messy Sequencing Data

**DOI:** 10.1101/281550

**Authors:** David Gerard, Luis Felipe Ventorim Ferrão, Antonio Augusto Franco Garcia, Matthew Stephens

## Abstract

Detecting and quantifying the differences in individual genomes (i.e. genotyping), plays a fundamental role in most modern bioinformatics pipelines. Many scientists now use reduced representation next-generation sequencing (NGS) approaches for genotyping. Genotyping diploid individuals using NGS is a well-studied field and similar methods for polyploid individuals are just emerging. However, there are many aspects of NGS data, particularly in polyploids, that remain unexplored by most methods. Our contributions in this paper are four-fold: (i) We draw attention to, and then model, common aspects of NGS data: sequencing error, allelic bias, overdispersion, and outlying observations. (ii) Many datasets feature related individuals, and so we use the structure of Mendelian segregation to build an empirical Bayes approach for genotyping polyploid individuals. (iii) We develop novel models to account for preferential pairing of chromosomes and harness these for genotyping. (iv) We derive oracle genotyping error rates that may be used for read depth suggestions. We assess the accuracy of our method in simulations and apply it to a dataset of hexaploid sweet potatoes (*Ipomoea batatas*). An R package implementing our method is available at https://cran.r-project.org/package=updog.

## 1 Introduction

New high-throughput genotyping methods [e.g. Davey et al., 2011] allow scientists to pursue important genetic, ecological and evolutionary questions for any organism, even those for which existing genomic resources are scarce [Chen et al., 2014]. These methods combine high-throughput sequencing with preparation of a reduced representation library, to sequence a small subset of the entire genome across many individuals. This strategy allows both genome-wide single nucleotide polymorphism (SNP) discovery and SNP genotyping at a reduced cost compared to whole-genome sequencing [Chen et al., 2014, Kim et al., 2016]. Specific examples of these methods include “restriction site-associated DNA sequencing” (RAD-seq) [Baird et al., 2008] and “genotyping-by-sequencing” (GBS) [Elshire et al., 2011]. Both of these approaches have been widely used in recent biological research, including in population-level analyses [Byrne et al., 2013, Schilling et al., 2014], quantitative trait loci mapping [Spindel et al., 2013], genomic prediction [Spindel et al., 2015], expression quantitative trait loci discovery [Liu et al., 2017], and genetic mapping studies [Shirasawa et al., 2017].

Statistical methods for SNP detection and genotype calling play a crucial role in these new genotyping technologies. And, indeed, considerable research has been performed to develop such methods [Nielsen et al., 2011]. Much of this research has focused on methods for diploid organisms — those with two copies of their genomes. Here, we focus on developing methods for polyploid organisms – specifically for autopolyploids, which are organisms with more than two copies of their genome of the same type and origin, and present polysomic inheritance [Garcia et al., 2013]. Autopolyploidy is a common feature in plants, including many important crops (e.g. sugarcane, potato, several forage crops, and some ornamental flowers). More generally, polyploidy plays a key role in plant evolution [Otto and Whitton, 2000, Soltis et al., 2014] and plant biodiversity [Soltis and Soltis, 2000], and understanding polyploidy is important when performing genomic selection and predicting important agronomic traits [Udall and Wendel, 2006]. Consequently there is strong interest in genotyping polyploid individuals, and indeed the last decade has seen considerable research into genotyping in both non-NGS data [Voorrips et al., 2011, Serang et al., 2012, Garcia et al., 2013, Bargary et al., 2014, Mollinari and Serang, 2015, Schmitz Carley et al., 2017] and NGS data [McKenna et al., 2010, Li, 2011, Garrison and Marth, 2012, Blischak et al., 2016, Maruki and Lynch, 2017, Blischak et al., 2018, Clark et al., 2018].

Here, we will demonstrate that current analysis methods, though carefully thought out, can be improved in several ways. Current methods fail to account for the fact that NGS data are inherently messy. Generally, samples are genotyped at low coverage to reduce cost [Glaubitz et al., 2014, Blischak et al., 2018], increasing variability. Errors in sequencing from the NGS platforms abound [Li et al., 2011] (Section 2.2). These are two well-known issues in NGS data. In this paper, we will further show that NGS data also face issues of systematic biases (e.g. resulting from the read-mapping step) (Section 2.3), added variability beyond the effects of low-coverage (Section 2.4), and the frequent occurrence of outlying observations (Section 2.5). Our first contribution in this paper is highlighting these issues on real data and then developing a method to account for them.

Our second contribution is to consider information from Mendelian segregation in NGS genotyping methods. Many experimental designs in plant breeding are derived from progeny test data [Li et al., 2014, Tennessen et al., 2014, McCallum et al., 2016, Shirasawa et al., 2017]. Such progenies often result from a bi-parental cross, including half and full-sib families, or from selfing. This naturally introduces a hierarchy that can be exploited to help genotype individuals with low coverage. Here, we implement this idea in the case of autopolyploids with polysomic inheritance and bivalent non-preferential pairing. Hierarchical modeling is a powerful statistical approach, and others have used its power in polyploid NGS SNP genotyping, not with Mendelian segregation, but in assuming Hardy-Weinberg equilibrium (HWE) or small deviations from HWE [Li, 2011, Garrison and Marth, 2012, Maruki and Lynch, 2017, Blischak et al., 2018]. Using Mendelian segregation for SNP genotyping has been used in non-NGS data [Serang et al., 2012, Schmitz Carley et al., 2017], and in diploid NGS data [Zhou and Whittemore, 2012], but to our knowledge we are the first to implement this for polyploid NGS data — though others have used deviations from Mendelian segregation as a way to filter SNPs [Chen et al., 2014, e.g.] or for the related problem of haplotype assembly [Motazedi et al., 2018].

Though wielding Mendelian segregation in species that exhibit bivalent non-preferential pairing can be a powerful approach, in some species this assumption can be inaccurate. Chromosomal hom(oe)ologues might exhibit partial or complete degrees of preferential pairing [Voorrips and Maliepaard, 2012]. Our next contribution is developing a model and inference procedures to account for arbitrary levels of preferential pairing in bivalent pairing species. In particular, this approach allows us to estimate if a species contains completely homologous chromosomes (autopolyploids), completely homoeologous chromosomes (allopolyploids), or some intermediate level of chromosomal pairing [Stift et al., 2008] [“segmental allopolyploids”, Stebbins, 1947]. We harness these estimates of preferential pairing to genotype (auto/allo)polyploids.

Our final contribution concerns the question raised by the development of our new model: “How many reads are needed to adequately genotype an individual?” This is a complicated by the fact that read depth calculations depend on many factors: the level of allelic bias, the sequencing error rate, the overdispersion, the distribution of genotypes in the sample, as well as the requirements of the intended downstream analyses. This last consideration is especially difficult to consider when coming up with general guidelines. For example, in association studies merely a strong correlation with the true genotype is necessary [Pritchard and Przeworski, 2001] and a smaller read depth might be adequate. Whereas for mapping studies, a few errors can inflate genetic maps [Hackett and Broadfoot, 2003] and so a larger read depth might be needed to improve accuracy. Rather than try to account for all of these contingencies, we took the approach of developing functions that a researcher may use to obtain lower bounds on the read depths required based on (i) the specifics of their data and (ii) the needs of their study. This is in alignment with the strategy in elementary statistics of not providing general guidelines for the sample size of a *t*-test, but rather provide functions to determine sample size based on the variance, effect size, power, and significance level.

Our paper is organized as follows. We develop our method using a real dataset [Shirasawa et al., 2017] as motivation in Section 2. During this development, we highlight several issues and solutions to genotyping NGS data, including overdispersion, allelic bias, and outlying observations. We then evaluate the performance of our method using Monte Carlo simulations (Sections 3.1 through 3.3) and demonstrate its superior genotyping accuracy to competing methods in the presence of overdispersion and allelic bias. We then use our method on a real dataset of hexaploid sweet potato (*Ipomoea batatas*) in Section 3.4. Oracle genotyping rates are explored in Sections 2.12 and 3.6. We finish with a discussion and future directions (Section 4).

## 2 Methods

We now describe models and methods for genotyping polyploid individuals. The models incorporate several features we have observed in real data. To help highlight these features, and for ease of explanation, we start with a simple model and gradually incorporate each additional feature. Notation is introduced gradually as required, and summarized for convenience in Table 1.

**Table 1:**
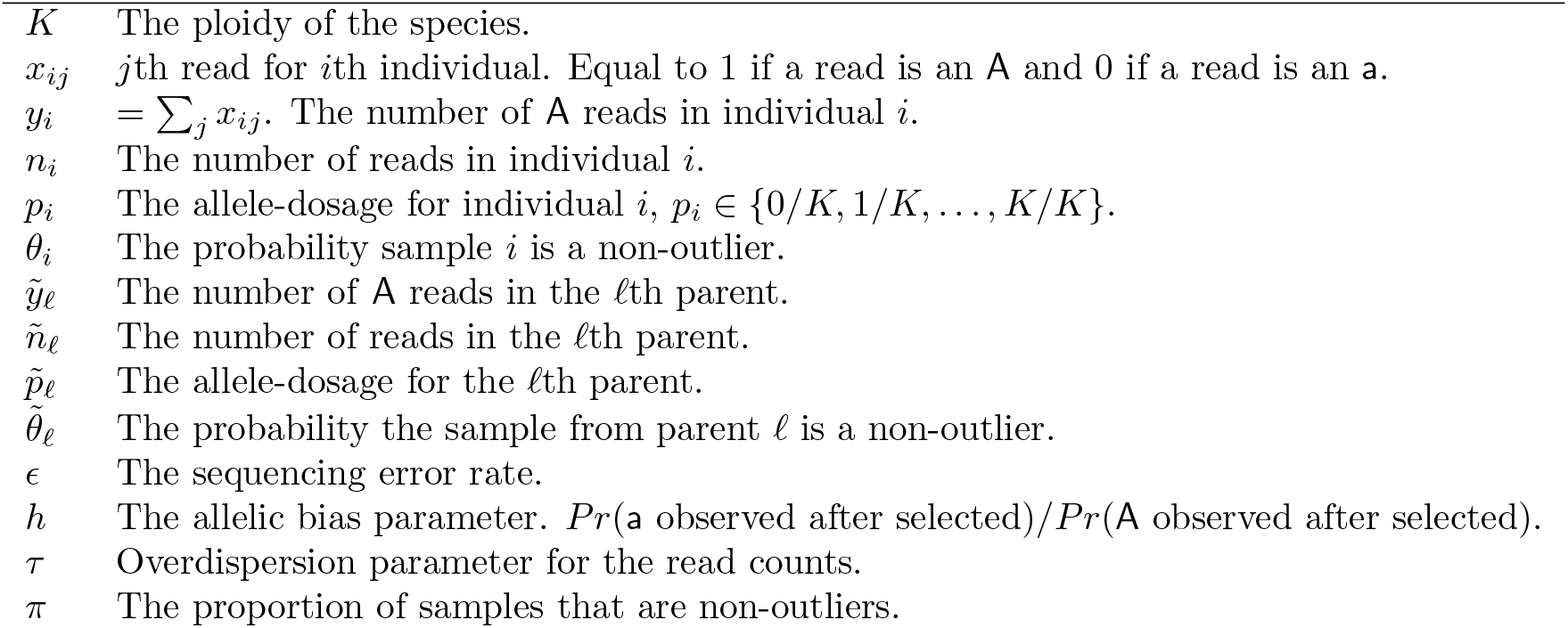
Summary of Notation.

To illustrate key features of our model we give examples from a dataset on autohexaploid sweet potato samples (*Ipomoea batatas*) (2n = 6x= 90) from a genetic mapping study by Shirasawa et al. [2017]. These data consist of an S1 population of 142 individuals, genotyped using double-digest RAD-seq technology [Peterson et al., 2012]. Here and throughout, “S1” refers to a population of individuals produced from the self-pollination of a single parent. We used the data resulting from the SNP selection and filtering procedures described in Shirasawa et al. [2017]. These procedures included mapping reads onto a reference genome of the related *Ipomoea trifida* to identify putative SNPs and then selecting high-confidence bi-allelic SNPs with coverage of at least 10 reads for each sample and with less than 25% of samples missing, yielding a total of 94,361 SNPs. Further details of the biological materials, assays, and data filtering may be found in Shirasawa et al. [2017].

For each SNP, we use A and a to denote the two alleles, with A being the reference allele (defined by the allele on the reference genome of the related *Ipomoea trifida*). For each individual the data at each SNP are summarized as the number of reads carrying the A allele and the number carrying the a allele. For a *K*-ploid individual there are *K* + 1 possible genotypes, corresponding to 0,1,…, *K* copies of the A allele. We use *p_i_* ∈ {0/*K*, 1/*K*,…, *K*/*K*} to denote the A allele dosage of individual *i* (so *Kp_i_* is the number of copies of allele A). Genotyping an individual corresponds to estimating *p_i_*.

Figure 1 illustrates the basic genotyping problem using data from a single well-behaved SNP. In this “genotype plot” each point is an individual, with the *x* and *y* axes showing the number of a and A reads respectively. The lines in the plot indicate the expected values for possible genotype *p_i_* ∈ {0/*K*, 1/*K*, …, *K*/*K*} and are defined by

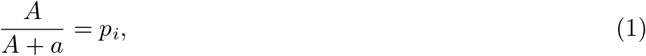

where *A* is the count of A reads and *a* is the count of A reads. Genotyping each sample effectively corresponds to determining which line gave rise to the sample’s data. In this case the determination is fairly clear because the SNP is particularly well-behaved and most samples have good coverage. Later we will see examples where the determination is harder.

**Figure 1:**
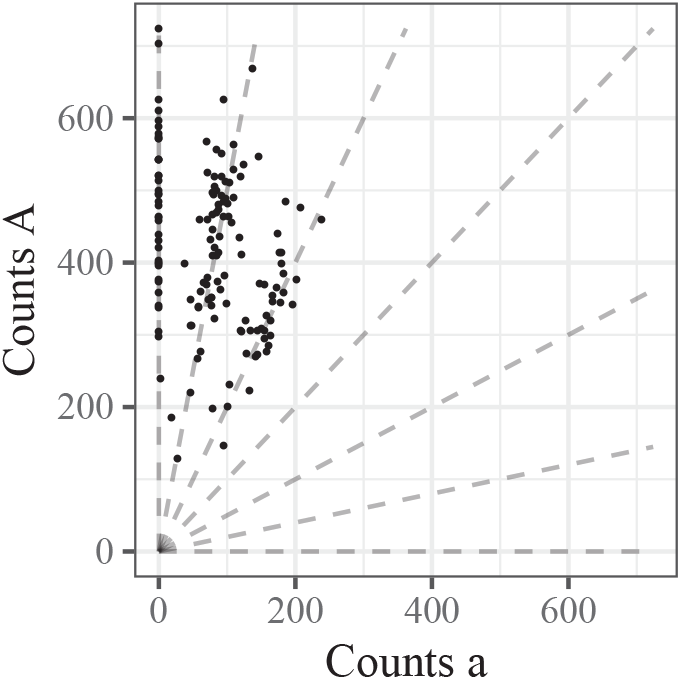
Genotype plot of single well-behaved SNP in a hexaploid species. Each point is an individual with the number of alternative reads along the *x*-axis and the number of reference reads along the *y*-axis. The dashed lines are defined by *y*/(*x* + *y*) = *p* for *p* ∈ {0/6,1/6,…, 6/6}, which correspond to genotypes {aaaaaa, Aaaaaa,…, AAAAAA}.

### 2.1 Naive model

A simple and natural model is that the reads at a given SNP are independent Bernoulli random variables:

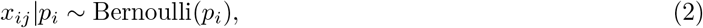

where *x_ij_* is 1 if read *j* from individual *i* is an A allele and is 0 if the read is an a allele. The total counts of allele A in individual i then follows a binomial distribution

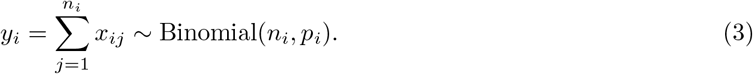

If the individuals are siblings then, by the rules of Mendelian segregation, the *p_i_*’s have distribution [Serang et al., 2012]:

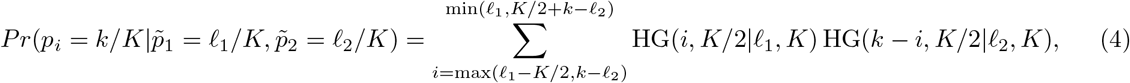

where 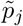 is the A allele dosage of parent *j* and HG(*a, b*|*c, d*) is the hypergeometric probability mass function:

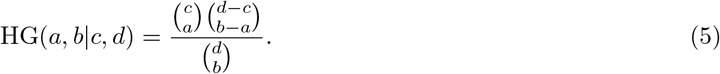

Equation (4) results from a convolution of two hypergeometric random variables. The distribution (4) effectively provides a prior distribution for *p_i_*, given the parental genotypes. If the parental genotypes are known then this prior is easily combined with the likelihood (3) to perform Bayesian inference for *p_i_*. If the parental genotypes are not known, then it is straightforward to estimate the parental genotypes by maximum likelihood, marginalizing out the *p_i_*, yielding an empirical Bayes procedure.

### 2.2 Modeling sequencing error

Though model (2) is a reasonable starting point, it does not account for sequencing error. Even if sequencing errors rates are low (e.g. 0.5-1% [Li et al., 2011]), it is crucial to model them because a single error can otherwise dramatically impact genotype calls. In particular, if an individual truly has all reference alleles (*p_i_* = 1) then (in the model without errors) a single non-reference allele observed in error would yield a likelihood (and hence posterior probability) of 0 for the true genotype. Biologically, this means that a homozygous individual can be erroneously classified as heterozygous, which may impact downstream analyses [Bourke et al., 2018]. We demonstrate this in Figure 2 where in the right panel we fit the model described in Section 2.1 to a single SNP. We labeled 6 points on Figure 2 with triangles that we think intuitively have a genotype of AAAAAA but were classified as having a genotype of AAAAAa due to the occurrence of one or two a reads.

**Figure 2:**
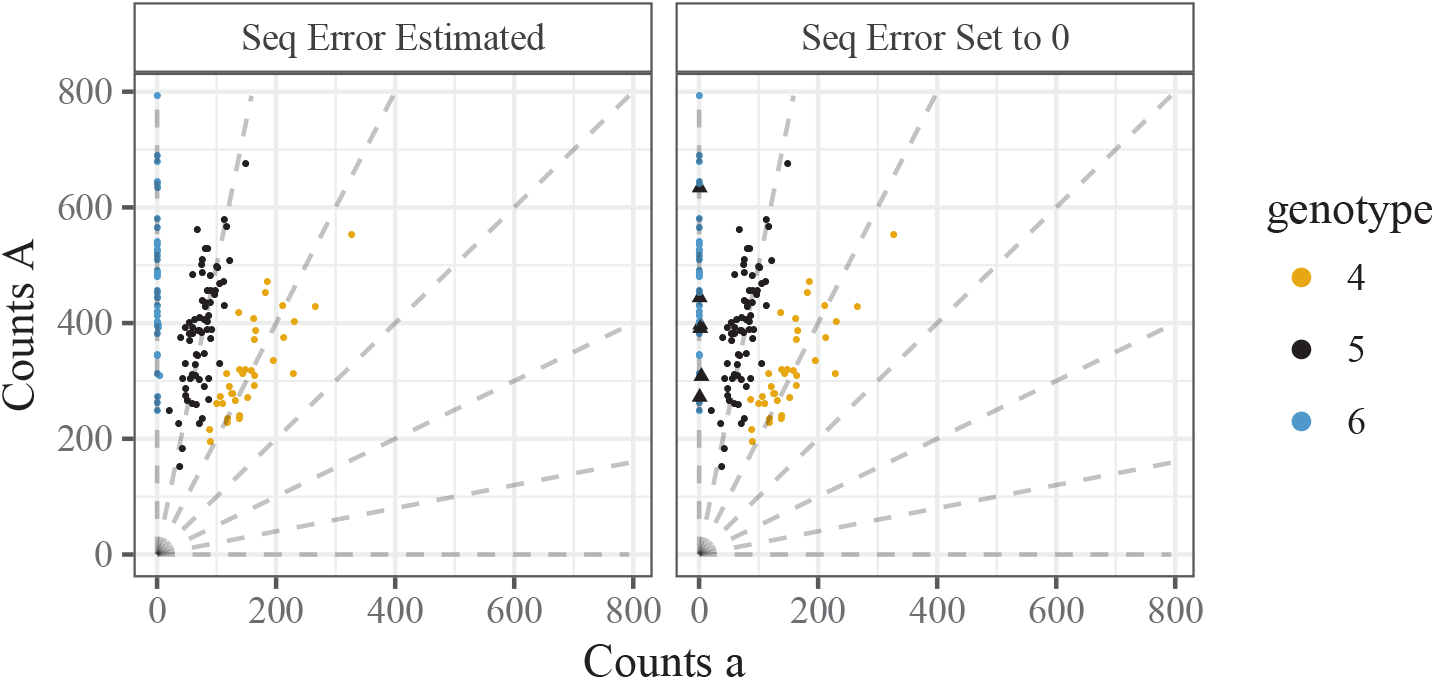
Two genotype plots demonstrating the need to model sequencing error. A SNP from hexaploid sweet potato genotyped in conjunction with estimating the sequencing error rate (left panel) or by setting the sequencing error rate to 0 (right panel). Triangles are points that we think look mis-classified.

To incorporate sequencing error, we replace (3) with

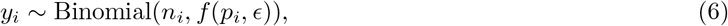

where

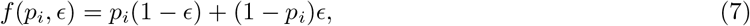

and *ϵ* denotes the sequencing error rate. Estimating this error rate in conjunction while fitting the model in Section 2.1 results in intuitive genotyping for the SNP in Figure 2 (left panel). This approach is also used by Li [2011] (and seems more principled than the alternative in Li et al. [2014]); see also Maruki and Lynch [2017] for extensions to multi-allelic SNPs.

### 2.3 Modeling allelic bias

We now address another common feature of these data: systematic bias towards one allele or the other. This is exemplified by the central panel of Figure 3. At first glance, it appears that all offspring have either 5 or 6 copies of the reference allele. However, this is unlikely to be the case: since this is an S1 population, if the parent had 5 copies of the reference allele, we would expect, under Mendelian segregation, the genotype proportions to be (0.25, 0.5, 0.25) for 6, 5, and 4 copies of the reference allele, respectively. Indeed, the proportion of individuals with greater than 95% of their read-counts being the reference allele is 0.197 — relatively close to the 0.25 expected proportion for genotype AAAAAA (one-sided *p*-value = 0.085). That leaves the other points to represent a mixture of AAAAAa and AAAAAa genotypes. Thus, for this SNP there appears to be bias toward observing an A read compared to an a read.

**Figure 3:**
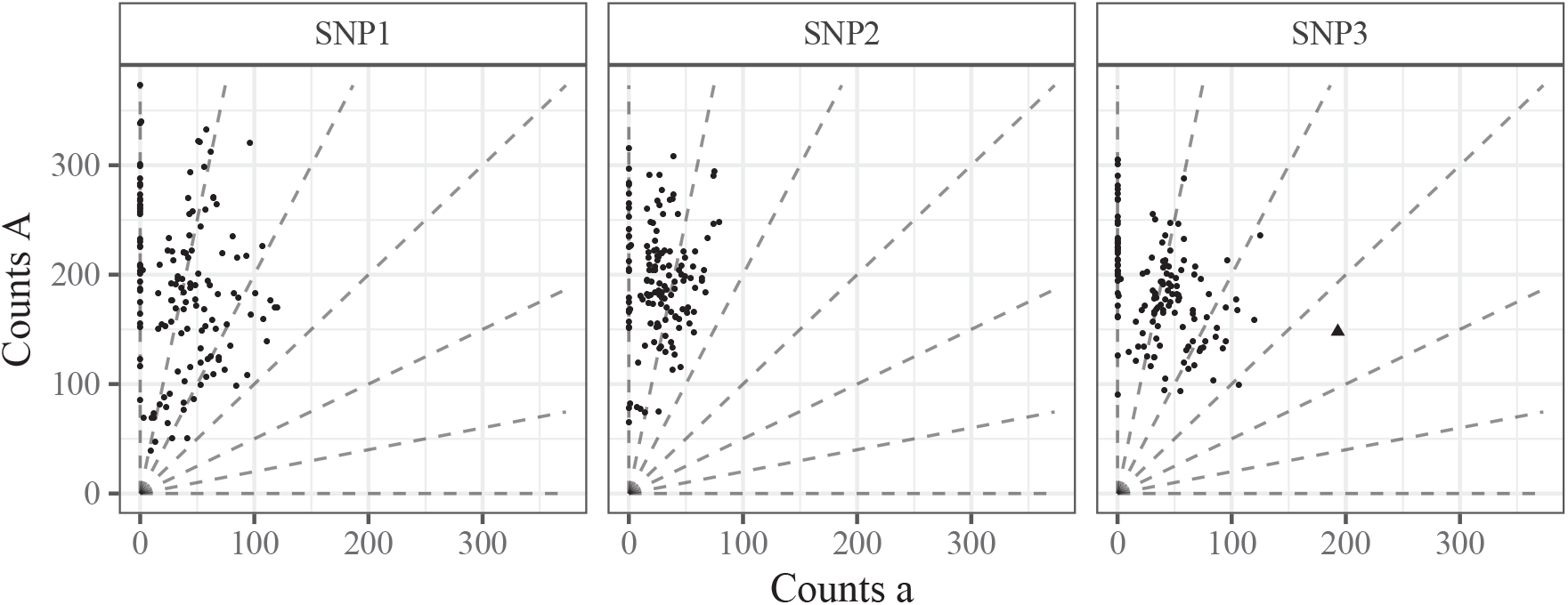
Three genotype plots of SNPs demonstrating common features of the data from Shirasawa et al. [2017], considering autohexaploid sweet potato: Overdispersion (left), bias (middle), and outlying observations (right).

One possible source of this bias is the read mapping step [Van De Geijn et al., 2015]. For example, if one allele provides a better match to a different location on the genome than the true location then this decreases its probability of being mapped correctly. Van De Geijn et al. [2015] describe a clever and simple technique to adjust for allele-specific bias during the read-mapping step. However, we see three possible problems that may be encountered in using the approach of Van De Geijn et al. [2015]: (i) in some instances, a researcher may not have access to the raw data files to perform this procedure; (ii) the procedure requires access to a reference genome, which is unavailable for many organisms [Lu et al., 2013]; (iii) there could plausibly be other sources of bias, requiring the development of a method agnostic to the source of bias.

To account for allelic bias, we model sequencing as a two stage procedure: first, reads are chosen to be sequenced (assumed independent of allele); and, second, chosen reads are either “observed” or “not observed” with probabilities that may depend on the allele they carry. Let *x_ij_* denote the random variable that is 1 if the chosen read carries an a allele, and 0 otherwise; and let *u_ij_* denote the random variable that is 1 if the chosen read is actually observed and 0 otherwise. To model the first stage we use

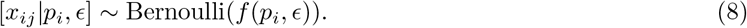

To model the second stage we assume

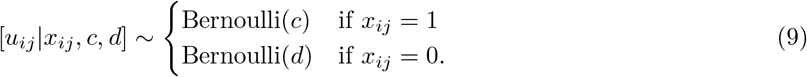

Allelic bias occurs when *c* ≠ *d*. Since we can only determine the alleles of the reads we observe, we are interested in the distribution of *x_ij_* conditioned on *u_ij_* = 1, which is given by Bayes rule:

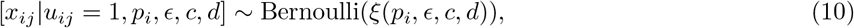

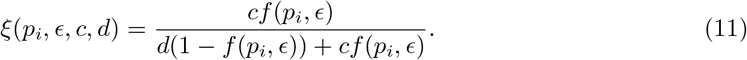

Notice that *ξ* depends on *c* and *d* only through the ratio *h* := *d/c*. Specifically:

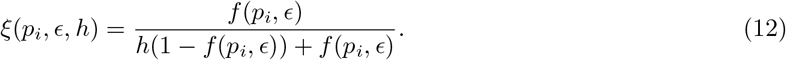

We refer to *h* as the “bias parameter”, which represents the relative probability of a read carrying the two different alleles being observed after being chosen to be sequenced. For example, a value of *h* = 1/2 means that an a read is twice as probable to be correctly observed than an a read, while a value of *h* = 2 means that an a read is twice as probable to be correctly observed than an a read.

Both the bias parameter *h* and the sequencing error rate *ϵ* modify the expected allele proportions for each genotype. However, they do so in different ways: lower values of *h* push the means toward the upper left of the genotype plot, while higher values push the means toward the lower right. Higher values of *ϵ* tend to squeeze the means toward each other. These different effects are illustrated in Figure 4.

**Figure 4:**
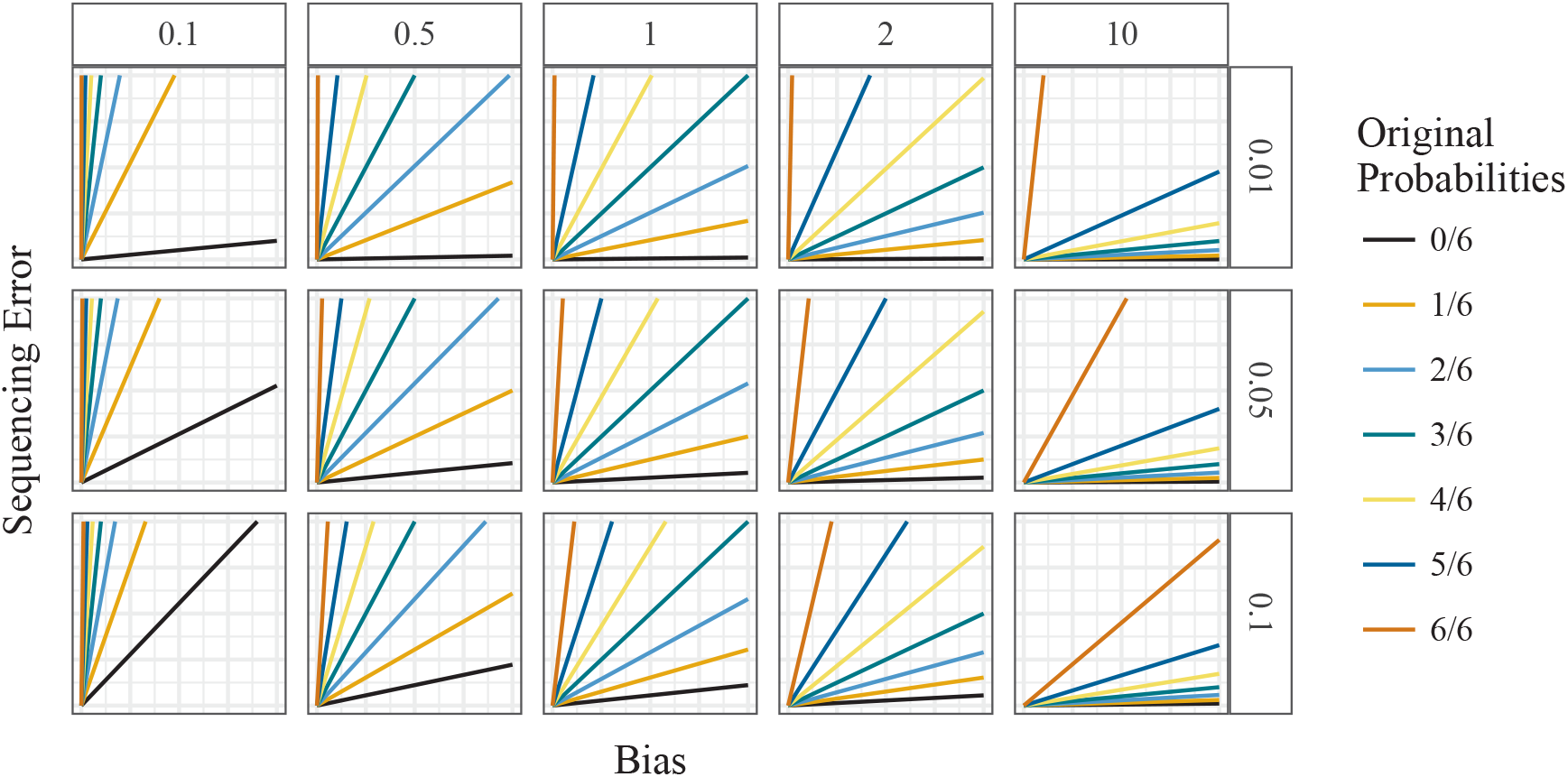
Mean counts of the mapped-reads under different levels of sequencing error rates (row facets) and different levels of allelic bias (column facets), considering an autohexaploid locus. The *y*-axis is the number of reference counts and the *x*-axis is the number of alternative counts.

### 2.4 Modeling overdispersion

Overdispersion refers to additional variability than expected under a simple model. Overdispersion is a common feature in many datasets. In sequencing experiments overdispersion could be introduced by variation in the processing of a sample, by variations in a sample’s biochemistry, or by added measurement error introduced by the sequencing machine. These could all result in observed read-counts being more dispersed than expected under the simple binomial model (3). Figure 5 (which is an annotated version of the left panel in Figure 3) illustrates overdispersion in these data.

**Figure 5:**
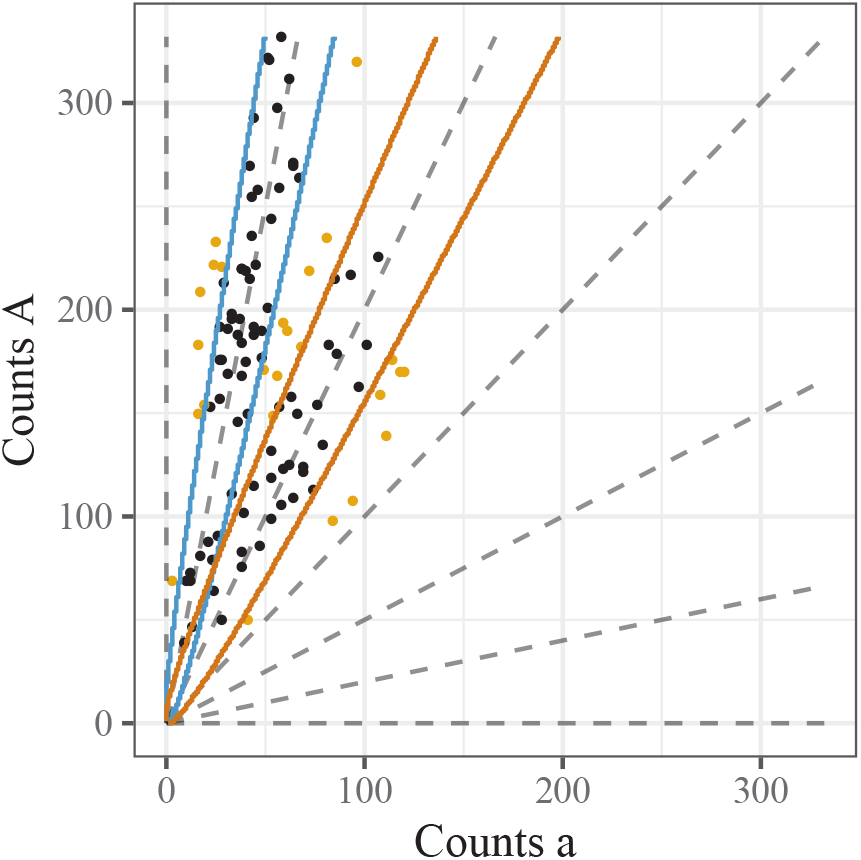
A genotype plot illustrating overdispersion compared with the simple binomial model. This figure shows the same SNP as the left panel of Figure 3 but with the points with greater than 95% a reads removed. Solid lines indicate the 0.025 and 0.975 quantiles for the Binomial distribution with probabilities 4/6 (red) and 5/6 (blue). Points that lie within these lines are colored black; outside are colored orange. Under the binomial model only 5% of the points should be orange, but in fact a significantly higher proportion (23.6%; *p*-value 1.2 × 10^−11^) are orange.

To model overdispersion we replace the binomial model with a beta-binomial model [Skellam, 1948]. The beta-binomial model assumes that each individual draws their own individual-specific mean probability from a beta-distribution, then draws their counts conditional on this probability:

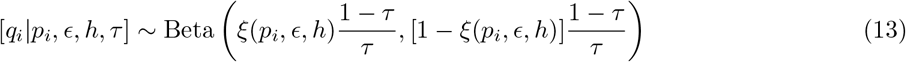

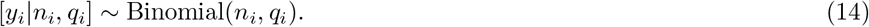

Here *ξ*(*p_i_, ϵ, h*) (12) is the mean of the underlying beta-distribution and *τ* ∈ [0,1] is the overdispersion parameter, with values closer to 0 indicating less overdispersion and values closer to 1 indicating greater overdispersion. (The parameter *τ* can also be interpreted as the “intra class correlation” [Crowder, 1979].) The *q_i_*’s are dummy variables representing individual specific probabilities and are integrated out in the following analyses. We denote the marginal distribution of *y_i_* as

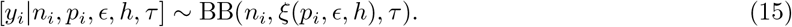

### 2.5 Modeling outliers

The right panel of Figure 3 illustrates another important feature of real data: outliers. Here, most of the points appear well-behaved, but one point (marked as a triangle) is far from the remaining points. Furthermore, taking account of the fact that these data came from an S1 population, this outlying point is inconsistent with the other points. This is because the other points strongly suggest that the parental genotype is AAAAAa, meaning that the possible genotypes are AAAAAA, AAAAAa, and AAAAAa, and the triangle lies far from the expectation for any of these genotypes. Indeed, under a fitted beta-binomial model, if the individual’s genotype were AAAAAa, then the probability of seeing as few or fewer a counts in this individual as were actually observed is 8.7 × 10^-6^ (Bonferroni-corrected *p*-value of 0.0012). There are many possible sources of outliers like this, including individual-specific quirks in the amplification or read-mapping steps, and sample contamination or mislabeling.

There are several common strategies for dealing with outliers. These include using methods that are inherently robust to outliers [Huber, 1964]; identifying and removing outliers prior to analysis [Hadi and Simonoff, 1993]; or modeling the outliers directly [Aitkin and Wilson, 1980]. Here we take this last approach, which has the advantage of providing measures of uncertainty for each point being an outlier.

Specifically, we model outliers using a mixture model:

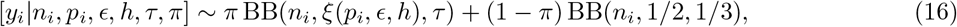

where *π* represents the proportion of points that are not outliers. Here the second component of this mixture represents the outliers, and the parameters (1/2,1/3) of this outlying distribution were chosen so that the underlying beta distribution is uniform on [0,1]. (We also tried estimating the underlying beta parameters for the outlying distribution, but we found that this can overfit the data in some instances; not shown.)

### 2.6 Prior on sequencing error rate and bias parameter

In some cases we found that maximum likelihood estimation of the bias parameter *h* and error rate *ϵ* gave estimates that were unrealistic, and lead to undesirable results. For example, we often observed this problem at SNPs where all individuals carry the same genotype (i.e. monomorphic markers).

Figure 6A shows an example of this problem for simulated data from an autotetraploid species in which an AAAA parent is crossed with an aaaa parent, which results in all offspring having the same genotype AAaa. Using the model described up to now yields a maximum likelihood estimate of sequencing error rate 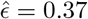, which is unrealistically high. This further creates very poor genotype calls (Figure 6B).

**Figure 6:**
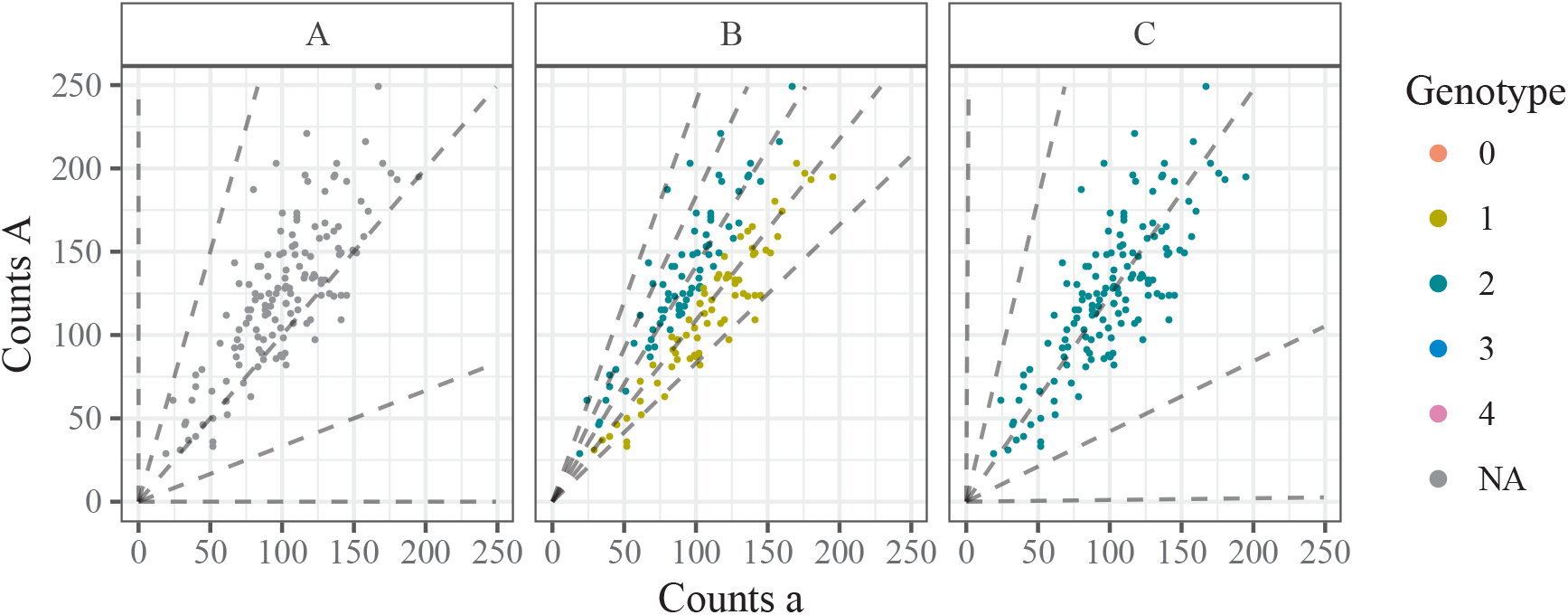
Three genotype plots illustrating the need to place priors on the bias and sequencing error rate parameters. Simulated autotetraploid NGS data where all individuals have the AAaa genotype. (A) unlabeled NGS data, (B) labeled by a fit without any penalties on the bias and sequencing error rate, and (C) labeled by a fit with penalties on both the bias and the sequencing error rate.

To avoid this problem we place priors on *ϵ* and *h* to capture the fact that e will usually be small and *h* will not deviate from 1 too far. Specifically we use

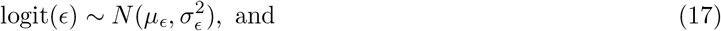

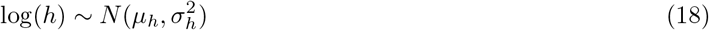

with software defaults *μ_ϵ_* = −4.7, 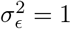, *μ_h_* = 0, and 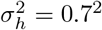. With these defaults, 95% of the prior mass is on *ϵ* ∈ [0.0012, 0.061] and *h* ∈ [0.25, 4.1] (see Figure S1 for graphical depictions.). We chose these defaults based on empirical observations (Figure S7), inflating the variances to make the priors more diffuse (less informative) than what we observe in practice. Our prior on the sequencing error rate is also consistent with literature estimates from various platforms [Goodwin et al., 2016], though some sequencing technologies can have error rates as high as 15% and, if using such technologies, the prior should be adjusted accordingly.

Ideally one would incorporate these prior distributions into a full Bayesian model, and integrate over the resulting posterior distribution on *ϵ* and *h*. However this would require non-trivial computation, and we take the simpler approach of simply multiplying the likelihood by the priors (17) and (18) and maximizing this product in place of the likelihood. (Effectively this corresponds to optimizing a penalized likelihood.) See Section 2.8 for details.

Using both these priors results in accurate genotypes for our simulated example (Figure 6C).

### 2.7 Incorporating parental reads

In genetic studies involving mating designs researchers almost always have NGS data on parent(s) as well as offspring. Such data can be easily incorporated into our model.

Specifically we model the number of a reads in parent *ℓ* (denoted 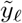) given the total number of reads in parent *ℓ* (denoted *ñ_ℓ_*) and allelic dosage *p_ℓ_* by:

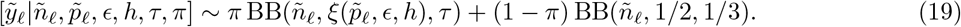

We treat the parental read count data as independent of the offspring count data (given the underlying genotypes), so the model likelihood is the product of (19) and (6).

### 2.8 Model

We now summarize our model and fitting procedure.

Our model is:

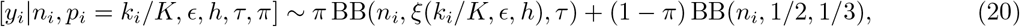

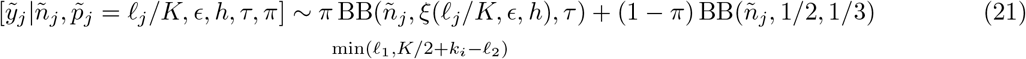

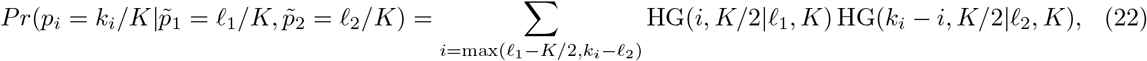

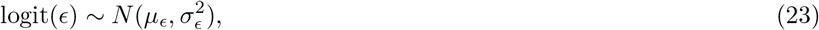

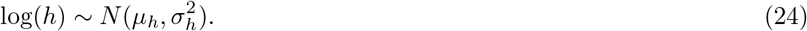

To fit this model for offspring from two shared parents (an F1 cross) we first estimate *ϵ, h, τ, π, ℓ*_1_, and *ℓ*_2_ via maximum likelihood (or, for *h, ϵ*, the posterior mode):

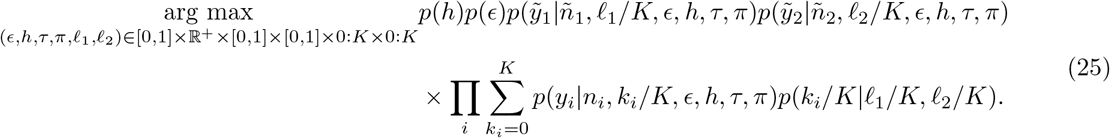

We perform this maximization using an expectation maximization (EM) algorithm, presented in Section A. The M-step (39) involves a quasi-Newton optimization for each possible combination of *ℓ*_γ_ and *ℓ*_2_, which we implement using the `optim` function in R [R Core Team, 2017].

Given estimates 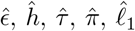, and 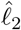, we use Bayes’ Theorem to obtain the posterior probability of the individuals’ genotypes:

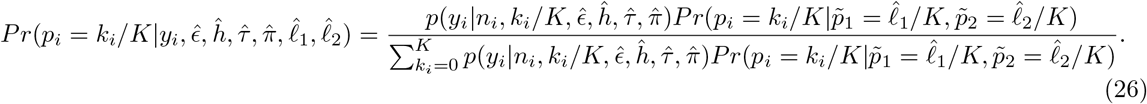

From this we can obtain, for example, the posterior mean genotype or a posterior mode genotype for each individual. The *θ_i_*’s from the E-step in (36) may be interpreted as the posterior probability that a point is a non-outlier.

Modifying the algorithm in Section A to deal with offspring from an S1 (instead of F1) cross is straightforward: simply constrain *ℓ*_γ_ = *ℓ*_2_ and remove 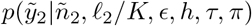 from (25).

This procedure is implemented in the R package updog (**U**sing **P**arental **D**ata for **O**ffspring **G**enotyping).

### 2.9 Screening SNPs

No matter the modeling decisions made, there will likely be poorly-behaved SNPs/individuals whose genotypes are unreliable. Such poorly behaved SNPs/individuals may originate from sequencing artifacts and it is important to consider identifying and removing them. The Bayesian paradigm we use naturally provides measures of genotyping quality, both at the individual and SNP levels.

Let *q_i_* be the maximum posterior probability of individual *i*. That is,

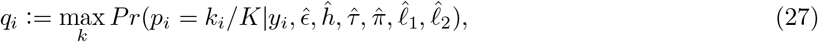

where the right-hand side of (27) is defined in (26). Then the posterior probability of an individual being genotyped incorrectly is 1 – *q_i_*, and individuals may be filtered based on this quantity. That is, if a researcher wants to have an error rate less than 0.05 then they would remove individuals with 1 – *q_i_* > 0.05. The overall error rate of a SNP is 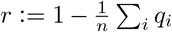, where *n* is the number of individuals in the sample. That is, *r* is the posterior proportion of individuals genotyped incorrectly at the SNP. Whole SNPs that have a large value of *r* may be discarded.

We evaluate the accuracy of *r* in Section 3.1.

### 2.10 Accounting for preferential pairing

Up until now, we have used a prior (4) that assumes a particular form of meiotic segregation — autopolyploids with polysomic inheritance and bivalent non-preferential pairing. If working with a species with different meiotic behavior — or a species where the meiotic behavior is unknown *a priori* [Bourke et al., 2017] — one should adjust this prior accordingly. In this section, we develop a model and inference procedures that account for one type of deviation from this ideal Mendelian segregation scenario: arbitrary levels of preferential pairing. Though, there are other ways to deviate from Mendelian segregation that we do not cover in this section. For example, if multivalent pairing were allowed then we would need to adjust our prior to account for the possibility of double reduction [Stift et al., 2010]. However, multivalent pairing can occur relatively infrequently in many stable autopolyploids [Bomblies et al., 2016], and when it does occur, the levels of double reduction can be relatively low [Stift et al., 2008], and so the below method might act as a reasonable approximation in the presence of multivalent pairing. Another example of Mendelian violations would be distortion resulting from an allele being semi-lethal.

Preferential pairing describes the phenomenon where some homologous chromosomes are more likely to pair during meiosis. Preferential pairing affects the segregation patterns of haplotypes and, hence, the distribution of genotypes in a sample. For a good overview of preferential pairing, see Voorrips and Maliepaard [2012].

To begin, we define a particular set of chromosome pairings to be a “pairing configuration”. Again, we suppose that all pairing during meiosis is bivalent. Since chromosomes are not labeled (we are only interested in the counts of reference and alternative alleles and not the specific chromosome carrying these alleles), we may represent each configuration by a 3-tuple 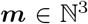 where the first element contains the number of aa pairs, the second element contains the number of Aa pairs, and the third element contains the number of AA pairs. For example, for the two configurations of a hexaploid parent with three copies of a,

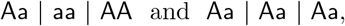

we represent the first configuration by *m* = (1,1,1) and the second by *m* = (0, 3, 0).

Given a configuration *m*, a *K*-ploid parent will segregate *K*/2 chromosomes to an offspring. The counts of reference alleles that segregate to the offspring, which we denote by *z*, will follow a “off-center” binomial distribution:

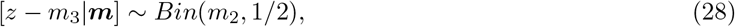

since in aa pairs no a alleles segregate, in AA pairs only a alleles segregate, and in Aa pairs an a allele segregates with probability 1/2. However, the configuration that forms ***m*** is a random variable. Suppose there are *q*(*ℓ*) possible configurations (see Theorem 1 for deriving *q*(*ℓ*)) given a parent has *I* copies of a. Also suppose configuration ***m**_i_* has probability *γ_i_* of forming. Then the distribution of *z* is a mixture of (28), with density

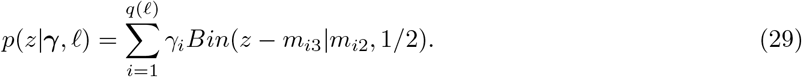

The values of the *γ_i_*’s in (29) determine the degree and strength of preferential pairing. If there is no preferential pairing, then the segregation probabilities (29) will equal those from the hypergeometric distribution (5). We derive the values of *γ_i_* that result in the hypergeometric distribution in Theorem 5. Any preferential pairing will result in deviations away from the weights in Theorem 5.

If neither parent in an F1 population exhibits preferential pairing, then the genotype distribution will follow (4). However, deviations from (4) can occur if some bivalent pairings occur with a higher probability than random chance would allow (Theorem 5). In which case, the offspring genotype distribution would be a convolution of two densities of family (29). Specifically, if parent 1 has genotype *ℓ*_1_ and configuration probabilities *γ*_1_ and parent 2 has genotype *ℓ*_2_ and configuration probabilities *γ*_2_, then the offspring genotype distribution is

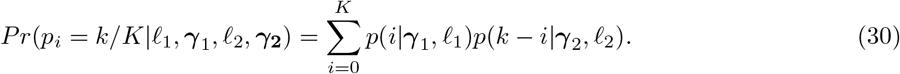

Allowing for arbitrary levels of preferential pairing is simply replacing prior (4) with (30). We have made this modification in updog, where we estimate both the parental genotypes (*ℓ*_1_ and *ℓ*_2_) and the degree and strength of the preferential pairing (*γ*_1_ and *γ*_2_) by maximum marginal likelihood.

This model for preferential pairing is similar to that of Stift et al. [2008]. Though there are major differences: (i) they do not use this model for genotyping, (ii) the chromosomes are not labeled in our setting and are in theirs, and (iii) we allow for arbitrary ploidy levels while they just allow for tetraploidy.

### 2.11 Extension to population studies

We have focused here on data from an F1 (or S1) experimental design, using parental information to improve genotype calls. However, a similar approach can also be applied to other samples (e.g. outbred populations) by replacing the prior on offspring genotypes (4) with another suitable prior. For example, previous studies have used a discrete uniform distribution [McKenna et al., 2010], a binomial distribution that results from assuming HWE [Li, 2011, Garrison and Marth, 2012], and a Balding-Nichols beta-binomial model on the genotypes [Balding and Nichols, 1995, 1997] that assumes an individual contains the same overdispersion parameter across loci [Blischak et al., 2018]. (All of these previous methods use models that are more limited than the one we present here: none of them account for allelic bias, outliers, or locus-specific overdispersion; and most implementations assume the sequencing error rate is known.)

In our software we have implemented both the uniform and binomial (HWE) priors on genotypes. The former is very straightforward. The latter involves modifying the algorithm in Section A by replacing *a_k_i__ℓ*_1_*ℓ*_2_ in (35) with the binomial probability *Pr*(*k_i_*|*n_i_, α*), where *α* is the allele frequency of A, and then optimizing over (*α,τ, h, ϵ*) in (39).

Though the uniform distribution might seem like a reasonable “non-informative” prior distribution, in practice it can result in unintuitive genotyping estimates. This is because the maximum marginal likelihood approach effectively estimates the allelic bias h and sequencing error rate e in such a way that the estimated genotypes approximate the assumed prior distribution. As most populations do not exhibit a uniform genotype distribution, this can result in extreme estimates of the bias and sequencing error rates, resulting in poor genotype estimates. We thus strongly recommend against using the uniform prior in practice and suggest researchers use more flexible prior distributions, such as the binomial prior.

### 2.12 Using oracle results for sampling depth guidelines

In this section, we develop oracle rates for incorrectly genotyping individuals based on our new likelihood (20). These rates are “oracle” in that they are the lowest achievable by an ideal estimator of genotypes. Using these oracle rates, we provide a method that returns a lower bound on read depths required to get an error rate less than some threshold. This method also returns the correlation of the oracle estimator with the true genotype, which might be useful when choosing the read depths for association studies.

To begin, we suppose that the sequencing error rate *ϵ*, bias *h*, overdispersion *τ*, and genotype distribution *π*(*k/K*) are all known. That is, the probability of dosage *p_i_* = *k/K* is *π*(*k/K*). Developing read depth suggestions when these quantities are known will provide lower bounds on the read depths required when they are not known. However, standard maximum likelihood theory guarantees that the following approximations will be accurate for a large enough sample of individuals (even for low read depth).

Given that these quantities are known, we have a simplified model with a likelihood *p*(*y_i_*|*p_i_* = *k*/*K*) (defined in (20)) and a prior *π*(*p_i_*), where the only unknown quantity is the allele dosage *p_i_*. Under 0/1 loss, the best estimator of *p_i_* is the maximum *a posteriori* (MAP) estimator

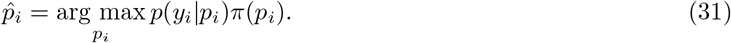

Notice that 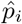 is only a function of *y_i_* and not *p_i_*. Thus, under this simplified model, one can derive the joint distribution of *p_i_* and 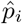 by summing over the possible counts *y_i_*.

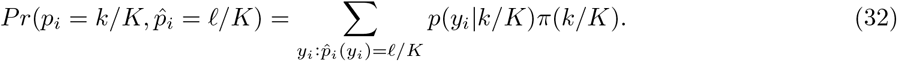

Given this joint distribution (32), one can calculate the oracle misclassification error rate

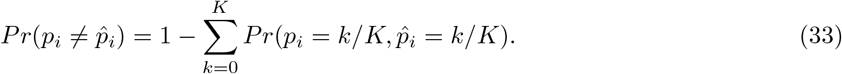

One can also calculate the oracle misclassification error rate conditioned on the individual’s genotype

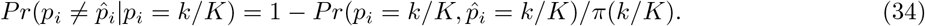

Also possible is calculating the correlation between the MAP estimator and the true genotype. All of these calculations have been implemented in the updog R package.

For sample size calculations, a researcher could specify the largest levels of bias, overdispersion, and sequencing error that they expect in their data. They could then specify a few genotype distributions that they expect (which is a finite number if the sample is from an F1 population), and then find the read depths required to obtain the oracle sequencing error rate (33) below some threshold that depends on their domain of application. We provide an example of this in Section 3.6.

In the machine learning community, equation (33) is often called the Bayes error rate [Hastie et al., 2009, Section 2.4].

### 2.13 Data availability

The methods implemented in this paper are available in the updog R package available on the Comprehensive R Archive Network at
https://cran.r-project.org/package=updog.

Code and instructions to reproduce all of the results of this paper (doi: 10.5281/zenodo.1203697) are available at
https://github.com/dcgerard/reproduce_genotyping.

## 3 Results

### 3.1 Simulation comparisons to other methods

We ran simulations to evaluate `updog’`s ability to estimate model parameters and genotype individuals in hexaploid species. Since most competing methods do not allow for an F1 population of individuals (though see Serang et al. [2012] implemented at https://bitbucket.org/orserang/supermassa.git), we compared the HWE version of `updog` (Section 2.11), a GATK-like method (from now on, just referred to as “GATK”) [McKenna et al., 2010] as implemented by `updog` (by using a discrete uniform prior on the genotypes, fixing *h* to 1 and *τ* to 0, while specifying *ϵ*), `fitPoly` [Voorrips et al., 2011], and the method of Li [2011] as implemented by Blischak et al. [2018]. Other methods are either specific to tetraploids, or only have implementations for tetraploids [Schmitz Carley et al., 2017, Maruki and Lynch, 2017], so we did not explore their performances.

Specifically, we simulated 142 unrelated individuals (the number of individuals in the dataset from Shirasawa et al. [2017]) under the `updog` model with sequencing error rate *ϵ* = 0.005, overdispersion parameter *τ* ∈ {0,0.01, 0.02}, and bias parameter *h* ∈ {0.5, 0.75,1} (with *h* = 1 indicating no bias). These parameter values were motivated by features in real data (Figure S7). We did not allow for outliers, neither in the simulated data nor in the fit. For each combination of *τ* and *h*, we simulated 1000 datasets. The *n_i_*’s were obtained from the 1000 SNPs in the Shirasawa et al. [2017] dataset with the largest read-counts. The distribution of the genotypes for each locus was binomially distributed using an allele frequency chosen from a uniform grid from 0.05 to 0.95.

Figure 7 contains boxplots for parameter estimates from the `updog` model. In general, the parameter estimates are highly accurate for small values of overdispersion and become less accurate, though still approximately unbiased, for larger values of overdispersion. The accuracy of the parameter estimates vary gradually for different levels of allele frequencies (Figures S3, S4, and S5). We note that in real data only a small fraction of estimates of the overdispersion parameter are higher than 0.02 (Figure S7).

**Figure 7:**
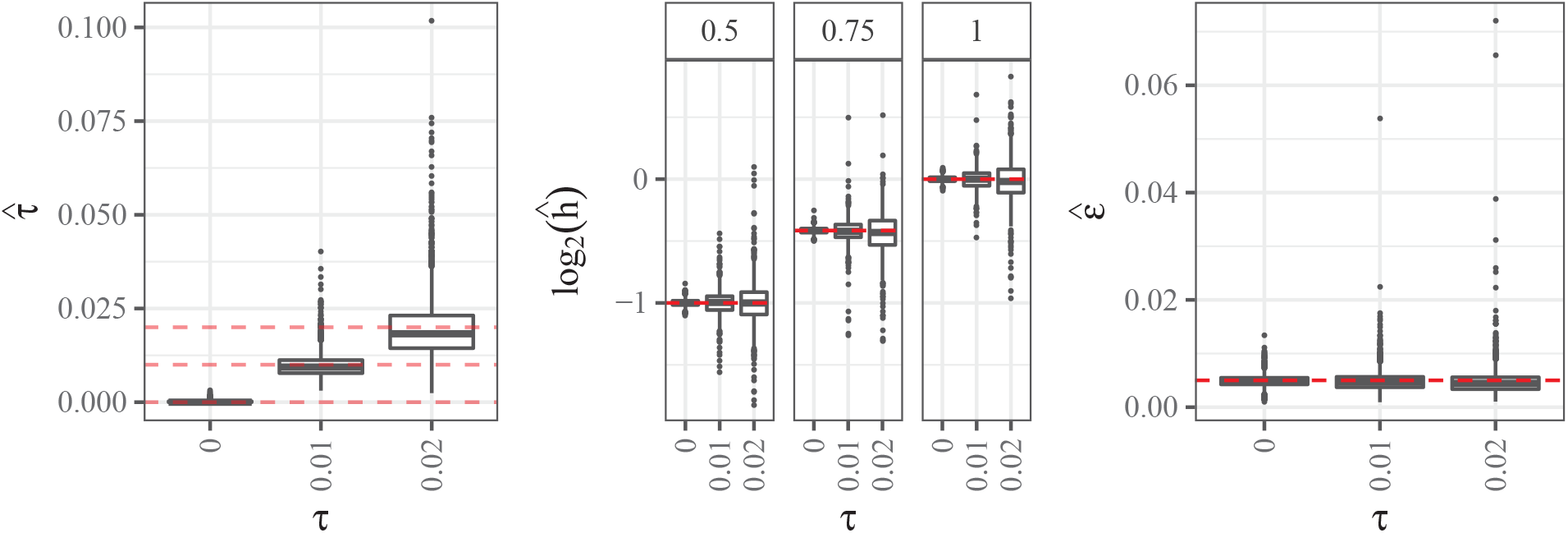
Left: Box plots of estimated overdispersion parameter (*y*-axis) stratified by the value of overdispersion (*x*-axis). The horizontal lines are at 0, 0.01, and 0.02. Center: Box plots of log_2_-transformed estimates of the bias parameter (*y*-axis) stratified by the value of overdispersion (*x*-axis). The column facets distinguish different levels of the true bias parameter (with 1 indicating no bias). The horizontal lines are at the true bias parameter. Right: Box plots of estimated sequencing error rate (*y*-axis) stratified by the value of overdispersion (*x*-axis). The horizontal line is at the true sequencing error rate of 0.005.

We then compared the genotyping accuracy of `updog` with the method from Li [2011], GATK [McKenna et al., 2010], and `fitPoly` [Voorrips et al., 2011]. In Figure 8 we have boxplots of the proportion of samples genotyped correctly stratified by bins of the allele frequency, color coding by method. We have the following conclusions:

1. All methods show similar performance when there is no bias (*h* = 1) and no overdispersion (*τ* = 0). These are the (unstated) assumptions of GATK and the method of Li [2011].
2. Overdispersion generally makes estimating genotypes much more difficult and accurate genotyping can only be guaranteed for small levels of overdispersion.
3. When there is any bias, `updog` has much superior performance to GATK and the method of Li [2011].
4. `fitPoly` performs well when there is no overdispersion, even for large levels of bias. However, in the presence of small amounts of overdispersion, `fitPoly` provides unstable genotype estimates.
5. Even when there is no bias, `updog` performs as well as the GATK and the method of Li [2011], except for some datasets where the allele frequency is close to 0.5 and the overdispersion is large. This is because GATK and the method of Li [2011] both effectively assume the bias parameter *h* is known at 1, while `updog` is using a degree of freedom to estimate the bias parameter. In the case where there is actually no bias, this can make `updog`’s estimates somewhat more unstable.

**Figure 8:**
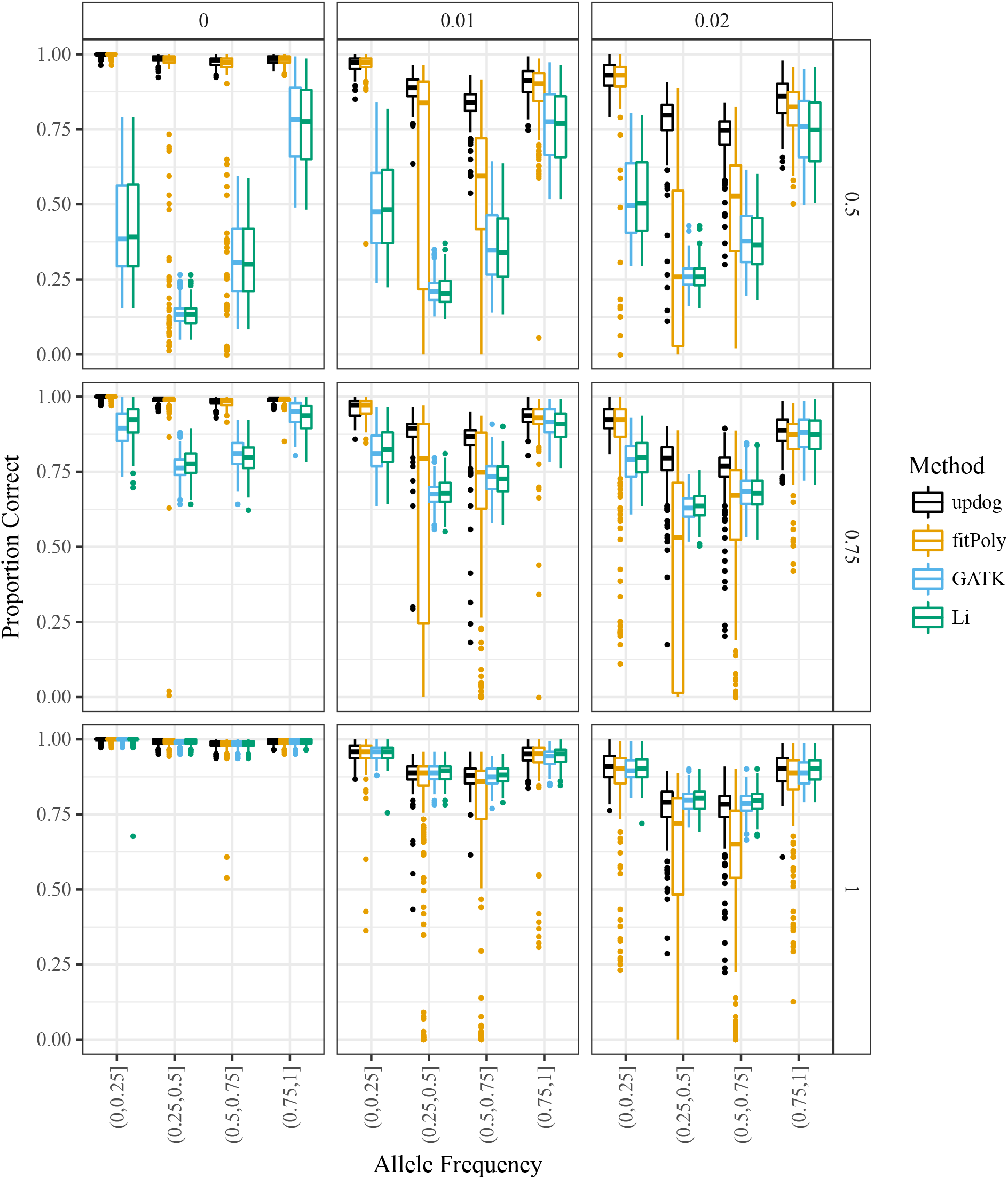
Boxplots of the proportion of individuals correctly genotyped (*y*-axis) for `updog` (black), `fitPoly` (orange), GATK (blue), and the method of Li [2011] (green) versus the true allele frequency (*x*-axis). The column facets distinguish between different levels of the overdispersion parameter and the row facets distinguish between different levels of the bias parameter (with 1 indicating no bias).

Even though the genotyping results are not encouraging for large amounts of overdispersion, `updog` has some ability to estimate this level of overdispersion to provide information to researchers on the quality of a SNP (Figure 7).

We compared the allele frequency estimates from `updog` and the method of Li [2011]. Here, the results are more encouraging (Figure S2). Even for large levels of overdispersion and bias, `updog` accurately estimates the allele frequency (though less accurately than with small overdispersion). As expected, the method of Li [2011] is adversely affected by bias, which it does not model, and tends to overestimate the allele frequency in the direction of the bias.

Also encouraging are the resulting correlations between the true and estimated genotypes. Figure S2 contains boxplots of the correlation of the true genotypes with the posterior means of each method — except for the method of Li [2011] where we calculate the correlation with the posterior mode as the software of Blischak et al. [2018] does not return posterior means. We used the posterior means because the posterior mode minimizes 0/1 loss, and so would be suboptimal for other, continuous metrics. We also included the naive method of returning the proportion of reference reads as the estimated genotype [Grandke et al., 2016]. From Figure S2, we see that `updog` outperforms all other methods in every scenario we explored. We see the largest gains over the naive method when there is no overdispersion and some bias, while the largest gains over `fitPoly` occur in high bias and high overdispersion scenarios. These results indicate that using `updog`’s posterior means could improve the results of association studies.

Finally, we compared the ability of `updog` and `fitPoly` to estimate the proportion of individuals genotyped incorrectly. Accurate estimates of this quantity can aid researchers when filtering for high-quality SNPs (Section 2.9). The software implementation of `updog` returns this quantity by default, while we derived this quantity from `fitPoly` using the output of the posterior probabilities on the genotypes. We subtracted the true proportion of individuals genotyped incorrectly from the estimated proportion and provide boxplots of these quantities in Figure 9. Generally, `updog` provides more accurate estimates for this proportion than `fitPoly`, particularly for small levels of overdispersion and allele frequencies close to 0.5.

**Figure 9:**
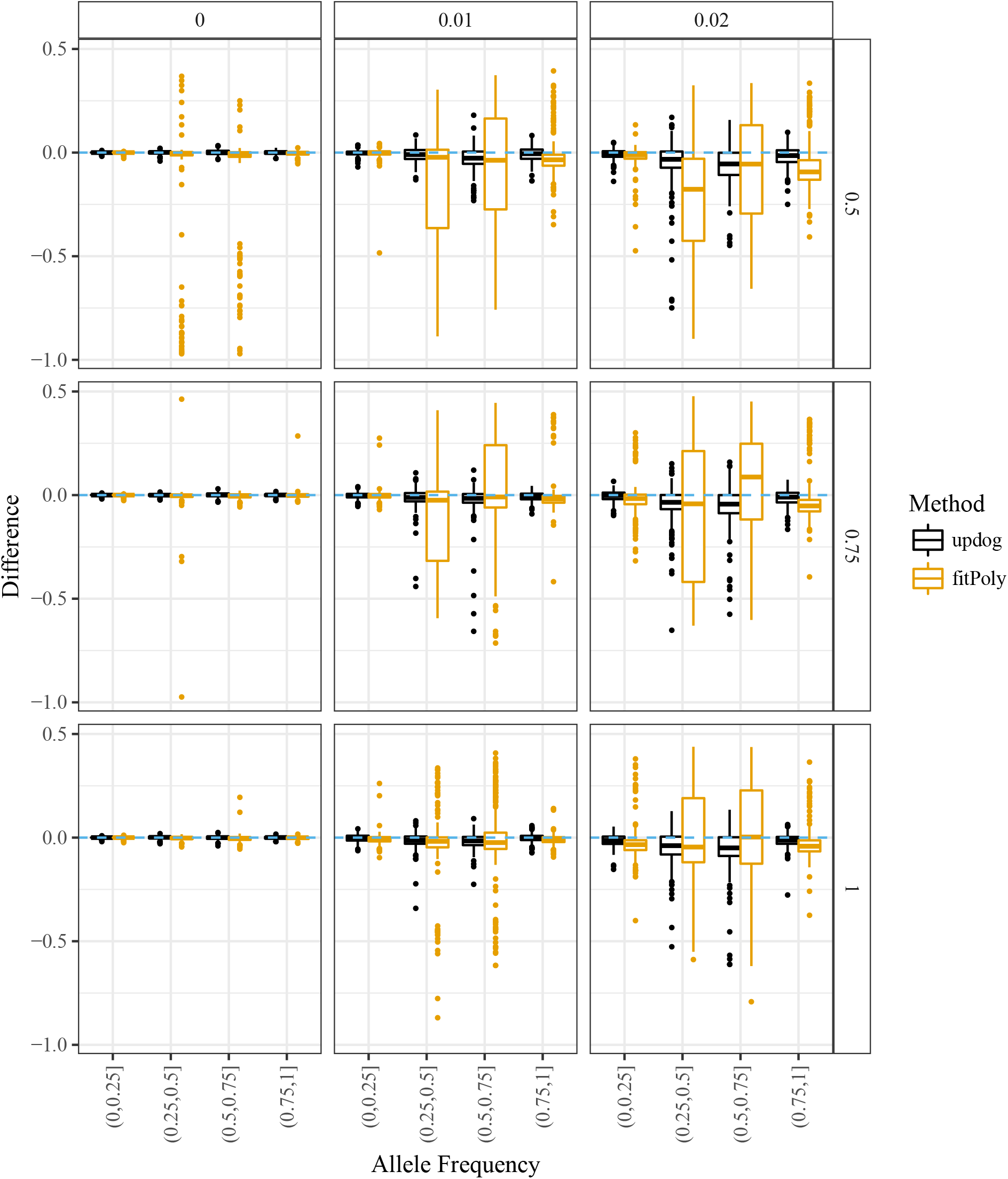
Difference between the estimated and true proportion of individuals genotyped incorrectly (*y*-axis) stratified by bins of the allele frequency (*x*-axis). Values above zero (the dashed horizontal line) correspond to overestimating the proportion of individuals genotyped incorrectly. The column facets distinguish between different levels of the overdispersion parameter and the row facets distinguish between different levels of the bias parameter (with 1 indicating no bias).

### 3.2 Simulation comparing use of S1 prior and HWE prior

We ran simulations to evaluate the gains in sharing information between siblings in an S1 population. We drew the genotypes of an S1 population of 142 individuals where the parent contains either 3, 4, or 5 copies of the reference allele. We then simulated these individuals’ read-counts under the `updog` model using the same parameter settings as in Section 3.1: *ϵ* = 0.005, *τ* ∈ {0,0.01,0.02}, and *h* ∈ {0.5, 0.75,1} (with *h* = 1 indicating no bias). For each combination of parental genotype, overdispersion, and allelic bias, we simulated 1000 datasets. The *n_i_*’s were again obtained from the 1000 SNPs in the Shirasawa et al. [2017] dataset with the largest read-counts.

For each dataset, we fit `updog` using a prior that either assumes the individuals were from an S1 population (Section 2.8) or were in HWE (Section 2.11). We plot summaries of the proportion of individuals genotyped correctly across datasets on Figure 10. In every combination of parameters, correctly assuming an S1 prior improves performance, particularly when there is a small amount of overdispersion (*τ* = 0.01). Box plots of the proportion of individuals genotyped correctly may be found in Figure S6.

**Figure 10:**
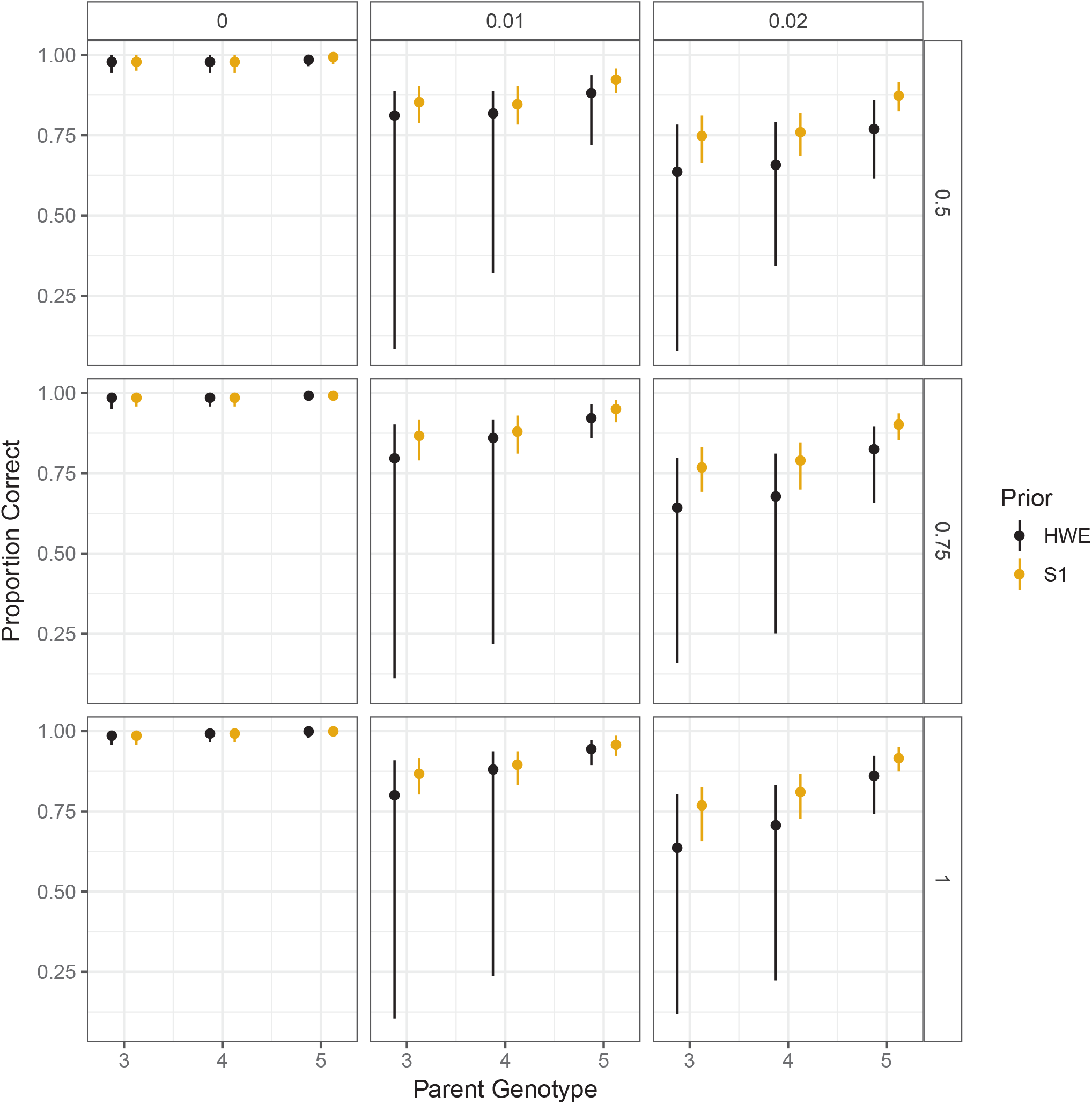
The 0.05, 0.5, and 0.95 quantiles (across datasets) of the proportion of individuals genotyped correctly in an `updog` fit (*y*-axis) stratified by parent genotype (*x*-axis). Column facets distinguish different levels of overdispersion while row facets distinguish different levels of bias (with 1 indicating no bias). Orange intervals correctly assume the individuals are from an S1 population while black intervals incorrectly assume the population is in Hardy-Weinberg equilibrium.

### 3.3 Simulations on preferential pairing

In Section 2.10 we developed a procedure to account for arbitrary levels of preferential pairing. In this section, we evaluate the accuracy of `updog` in the presence of preferential pairing.

We consider a tetraploid S1 population of individuals. In such a population, preferential pairing only affects the segregation probabilities when the parent has two copies of the reference allele (because this is the only scenario in a tetraploid population where there is more than one pairing configuration — see Theorem 1). In this setting, the two possible pairing configurations are ***m*** = (1,0,1) and ***m*** = (0, 2, 0). Under a non-preferential pairing setting, the weights of each configuration would be *γ*_1_ = 1/3 for the first configuration and *γ*_2_ = 2/3 for the second (Theorem 5).

To explore the performance of `updog` in the case of the most extreme levels of preferential pairing possible in a tetraploid species, we simulated offspring genotypes by setting either *γ*_1_ = 0 or *γ*_1_ = 1. We did this while setting *ϵ* = 0.005, *τ* ∈ {0,0.01, 0.02}, and *h* ∈ {0.5, 0.75,1}. Figure 11 contains boxplots of the estimated levels of *γ*_1_ against the true values of *γ*_1_. `updog` can estimate *γ*_1_ reasonably accurately, though these estimates become less stable as the overdispersion increases.

**Figure 11:**
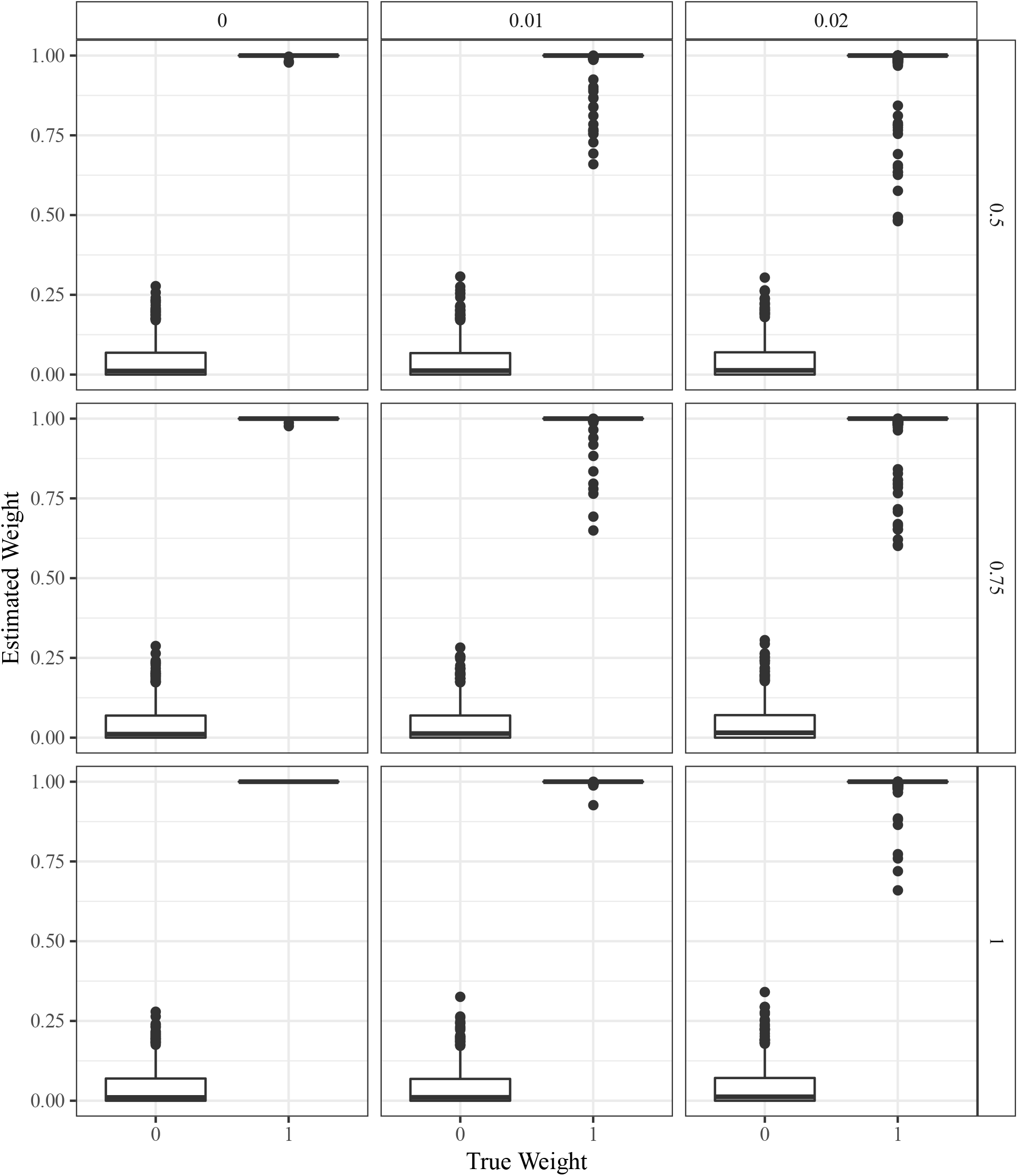
Boxplots of the estimated mixing proportion of one of the two possible pairing configurations (*y*-axis) stratified by the actual mixing proportion (*x*-axis) in a tetraploid S1 population where the parent has two copies of the reference allele. The row facets index the allelic bias (where 1 indicates no bias) and the column facets index the overdispersion.

We compared the preferential-pairing version of `updog` to `fitPoly` and the non-preferential pairing version of `updog`. Figure 12 contains boxplots of the proportion of individuals genotyped correctly stratified by *γ*_1_. The preferential pairing version of `updog` performs better than `fitPoly` and the non-preferential pairing version of `updog`, particularly in the presence of large amounts of overdispersion, and when *γ*_1_ is large.

**Figure 12:**
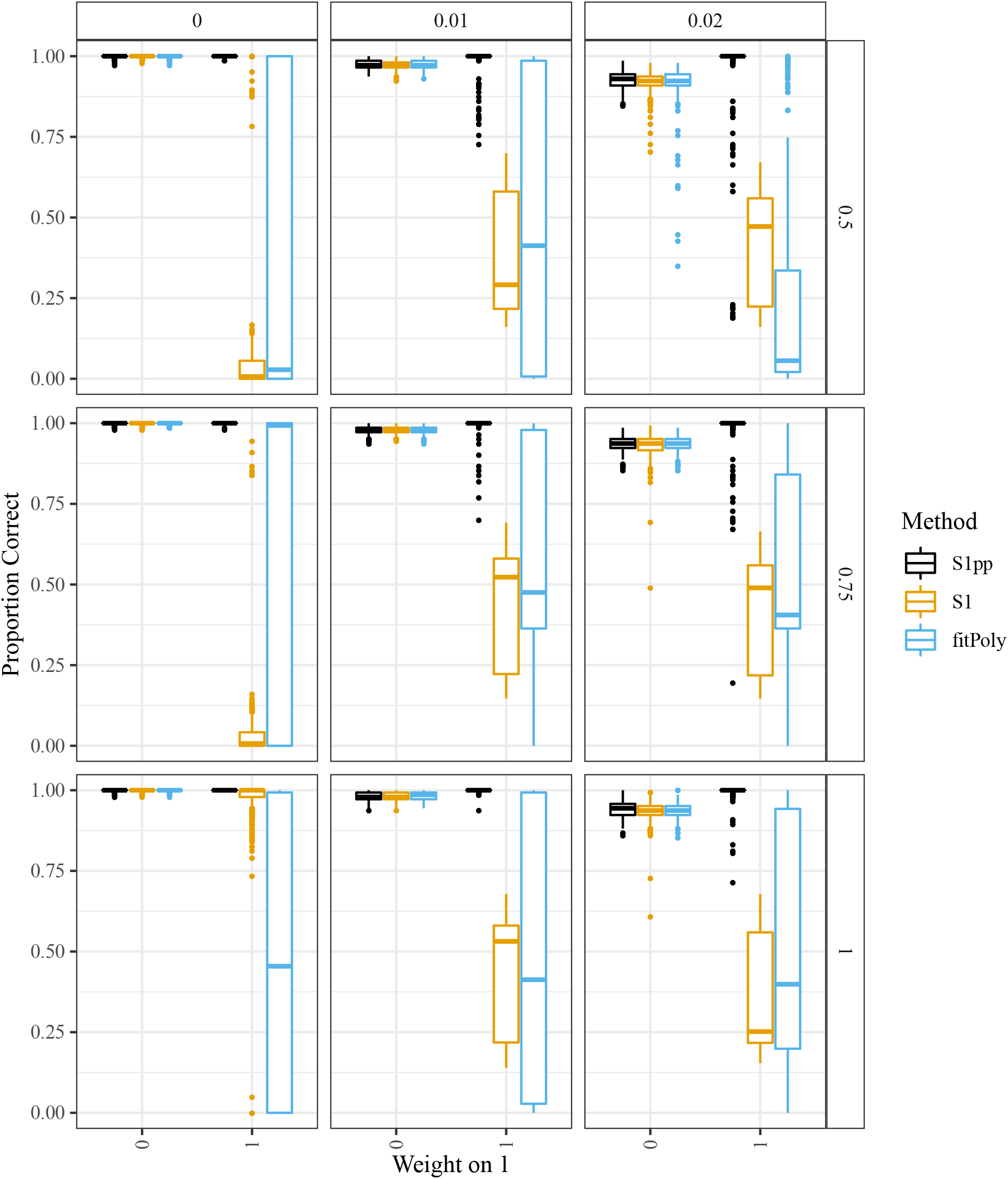
Boxplots of the proportion of individuals genotyped correctly (*y*-axis) stratified by the weight on one of the two possible pairing configurations (*x*-axis) in a tetraploid S1 population where the parent has two copies of the reference allele. The row facets index the allelic bias (where 1 indicates no bias) and the column facets index the overdispersion. Boxplots are color-coded by method: an `updog` fit that accounts for preferential pairing (black), an `updog` fit that does not account for preferential pairing (orange), and `fitPoly` (blue).

### 3.4 Sweet potato

We fit the Balding-Nichols version of the method of Blischak et al. [2018], the method of Serang et al. [2012] (implemented in the SuperMASSA software at http://statgen.esalq.usp.br/SuperMASSA/), `fitPoly` [Voorrips et al., 2011], and `updog` to the SNPs presented in Figure 3. The Balding-Nichols model that Blischak et al. [2018] uses has parameters that are shared across loci to estimate the genotype distributions of the SNPs, so we actually fit the method of Blischak et al. [2018] on the 1000 SNPs with the most read-counts and just present the results for the three SNPs from Figure 3. As the method of Blischak et al. [2018] requires the sequencing error rate to be known, we use the sequencing error rate estimates provided by `updog`.

**Figure 13:**
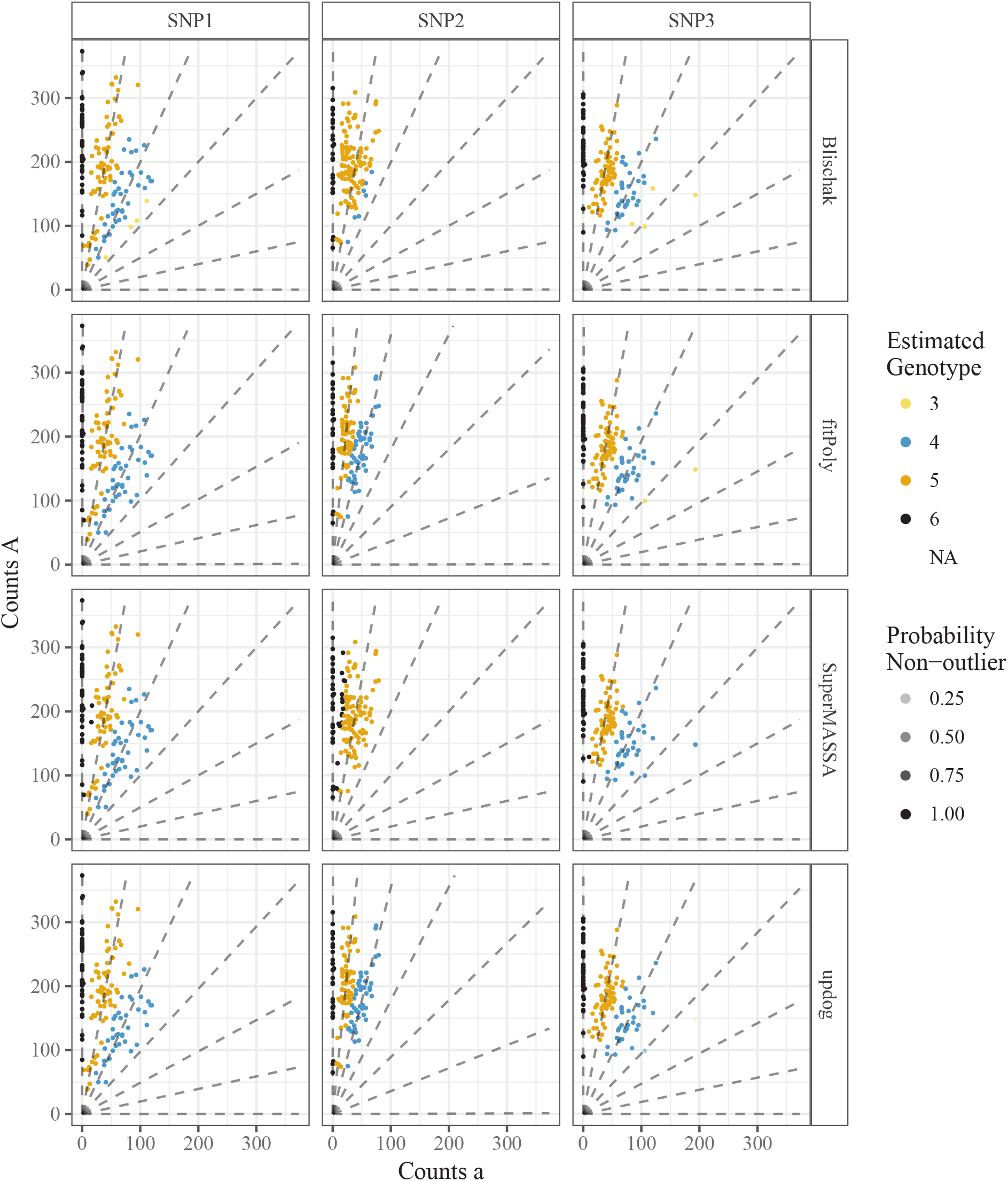
Genotype plots as in Figure 3 but color-coded by the estimated genotypes from the method of Blischak et al. [2018], the method of Serang et al. [2012], `fitPoly` [Voorrips et al., 2011], or `updog` (column facets). The column facets distinguish between the different SNPs. Transparency is proportional to the probability of being an outlier (only for `updog`).

`FitPoly` allows for F1, but not S1, populations, and seemingly only when parental counts are provided. To account for this, we copied the read-counts data from the parent and specified this duplicated point as a second parent. As recommended by the authors’ documentation, we fit these data using the `try.HW = TRUE` setting.

The fits for all four methods can be seen in Figure 13. Our conclusions are:

1. Serang et al. [2012] and Blischak et al. [2018] provide unintuitive results for SNP2. As we know this is an S1 population, the genotype distribution should be closer to a 1:2:1 ratio for genotypes AAAAAA, AAAAAa, and AAAAAa. Both `updog` and `fitPoly` correctly accounts for this. `Updog` does so by estimating an extreme bias while `fitPoly` does so by allowing the mean proportion of reference counts for each genotype to depend only linearly or quadratically on the dosage level.
2. The method of Serang et al. [2012] was designed for Gaussian, not count, data and as such provides some unintuitive genotyping. In particular, for SNPs 1 and 2 we see a few points on the left end of the AAAAAa genotypes that are coded AAAAAa when these could probably only result from an extreme sequencing error rate given that genotype.
3. The point in SNP3 that we have described as an outlier is removed by `updog`. The other methods are not able to cope with outliers.

Generally, `fitPoly` returns more similar genotyping estimates to `updog` on these data than other methods. However, their posterior measures of uncertainty differ, and sometimes substantially. Figure S8 plots the estimates for the proportion of individuals genotyped incorrectly by both `updog` and `fitPoly`. `Updog` usually returns larger levels of uncertainty, though there are some datasets where `fitPoly` estimates that a much larger proportion of individuals are genotyped incorrectly than `updog`. To us, these seem like anomalous results by `fitPoly`, as these SNPs seem mostly well-behaved (Figure S9).

### 3.5 Computation time

We measured the computation time required to fit `updog` to the 1000 SNPs with the highest read depth from the dataset of Shirasawa et al. [2017]. These computations were run on a 4.0 GHz quad-core PC running Linux with 32 GB of memory. It took on average 3.3 seconds per SNP to fit `updog`, with 95% of the runs taking between 1.81 and 6.56 seconds. This is much larger than the total time of 121 seconds it took for the software of Blischak et al. [2018] to fit their model on all 1000 SNPs. But it is almost exactly equal to the computation time of `fitPoly`, where 95% of the runs took between 1.77 and 6.55 seconds. Also, since the SNPs in `updog` are fitted independently, this allows us to easily parallelize this computation over the SNPs.

### 3.6 Sampling depth suggestions

In Section 2.12, we derived the oracle genotyping error rates given that the sequencing error rate, the allelic bias, the overdispersion, and the genotype distributions are all known. In this section, we explore these functions in the particular case when a SNP is in HWE with an allele frequency of 0.8.

We found the minimum read depths needed to obtain an error rate (33) less than 0.05 under different levels of ploidy *K* ∈ {2,4, 6}, bias *h* ∈ {0.5, 0.55, 0.6,…, 1}, and overdispersion *τ* ∈ {0, 0.002, 0.004,…, 0.02} while fixing the sequencing error rate at *ϵ* = 0.001. A heatmap of the minimum read depths required for the different ploidy levels may be found in Figures S10-S12. We have the following conclusions: (i) For diploid individuals, the levels of bias and overdispersion matter relatively little in terms of genotyping accuracy, and accurate genotyping may be made with an extremely small read depth. (ii) Read depth requirements seem to grow exponentially with the ploidy. (iii) The levels of bias and overdispersion greatly impact the read depth requirements in higher ploidy species. For reasonable levels of bias and overdispersion (Figure S7), the read depth requirements can be many times larger to obtain the same error rates as when there is no bias and no overdispersion.

Though the read depth requirements to obtain a small error rate seem depressing under reasonable levels of bias and overdispersion, the results are much more optimistic to obtain high correlation with the true genotypes. In Figures S13-S15, we calculated the minimum read depths required to obtain a correlation with the true genotypes of greater than 0.9 under the same parameter settings as Figures S10-S12. There, we see that the read depth requirements are much more reasonable, where for hexaploid species a read depth of 90 is adequate to obtain this correlation under large levels of bias and overdispersion. Correspondingly, for tetraploids a read depth of 25 seems adequate to obtain high correlation under large levels of bias and overdispersion.

Once again, we would like to stress that these are *lower bounds* on the read depth requirements. Actual read depths should be slightly larger than these, based on how many individuals are in the study.

## 4 Discussion

We have developed an empirical Bayes genotyping procedure that takes into account common aspects of NGS data: sequencing error, allelic bias, overdispersion, and outlying points. We have shown that accounting for allelic bias is vital for accurate genotyping, and that that our posterior measures of uncertainty (automatically taking into account the estimated levels of overdispersion) are well calibrated and may be used as quality metrics for SNPs/individuals. We confirmed the validity of our method on simulated and real data.

We have focused on a dataset that has a relatively large read-coverage and contains a large amount of known structure (via Mendelian segregation). In many datasets, one would expect to have much lower coverage of SNPs and less structure [Blischak et al., 2018]. For such data, we do not expect the problems of allelic bias, overdispersion, and outlying points to disappear. From our simulations, the most insidious of these issues to ignore is the allelic bias. If a reference genome is available, then it might be possible to correct for the read-mapping bias by using the methods from Van De Geijn et al. [2015]. However, this might not be the total cause of allelic bias. And without a reference genome, it is important to model this bias directly. We do not expect small coverage SNPs to contain enough information to accurately estimate the bias using our methods. For such SNPs, more work is needed. It might be possible to borrow strength between SNPs to develop accurate genotyping methods.

We have assumed that the ploidy is known and constant between individuals. However, some species (e.g. sugarcane) can have different ploidies per individual [Garcia et al., 2013]. If one has access to good cytological information on the ploidy of each individual, it would not be conceptually difficult to modify `updog` to allow for different (and known) ploidies of the individuals. However, estimating the ploidy might be more difficult, particularly in the presence of allelic bias. In the presence of such bias, one can imagine that it would be difficult to discern if a sample’s location on a genotype plot was due to bias or due to a higher or lower ploidy level. More work would be needed to develop an approach that works with individuals having unknown ploidy levels. Serang et al. [2012] attempts to estimate the ploidy level in Gaussian data, but they do not jointly account for allelic bias, which we hypothesize would bias their genotyping results.

Garrison and Marth [2012] uses a multinomial likelihood to model multiallelic haplotypes. This allows for more complex genotyping beyond SNPs. The models presented here could be easily extended to a multinomial likelihood. For example, to model bias with k possible alleles, one could introduce *k* – 1 bias parameters which measure the bias of each allele relative to a reference allele. One could use a Dirichlet-multinomial distribution to model overdispersion and a uniform-multinomial distribution (uniform on the standard *k* – 1 simplex) to model outliers. Modeling allele-detection errors (corresponding to sequencing errors in our NGS setup) would be context-specific.

The methods developed here may also be useful for genotyping diploid individuals. We are not aware of diploid genotyping methods that account for allelic bias, overdispersion, and outlying points. However, considering our oracle results in Section 3.6, accounting for these features is probably more important for polyploid species, as determining allele dosage is more difficult than determining heterozygosity/homozygosity.

Genotyping is typically only one part of a large analysis pipeline. It is known that genotyping errors can inflate genetic maps [Hackett and Broadfoot, 2003], but it remains to determine the impact of dosage estimates on other downstream analyses. In principle one would want to integrate out uncertainty in estimated dosages, but in diploid analyses it is much more common to ignore uncertainty in genotypes and simply use the posterior mean genotype — for example, in GWAS analyses [Guan and Stephens, 2008]. It may be that similar ideas could work well for GWAS in polyploid species (see also Grandke et al. [2016]).

## 5 Acknowledgments

We sincerely thank the authors of Shirasawa et al. [2017] for providing their data and Paul Blischak for providing useful comments. D. Gerard and M. Stephens were supported by NIH grant HG002585 and by a grant from the Gordon and Betty Moore Foundation (Grant GBMF #4559). L. F. V. Ferrao and A.A.F. Garcia were partially supported by grant 2014/20389-2, FAPESP/CAPES (São Paulo Research Foundation). A.A.F Garcia was supported by a productivity scholarship from the National Council for Scientific and Technological Development (CNPq).

## A EM algorithm for an F1 cross

1. Let (see (4))

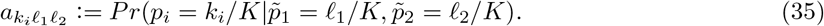
2. E-Step: Set

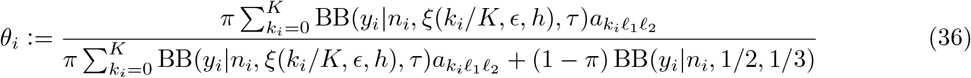

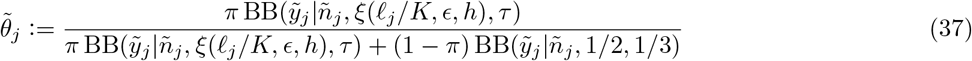
3. M-Step: Set

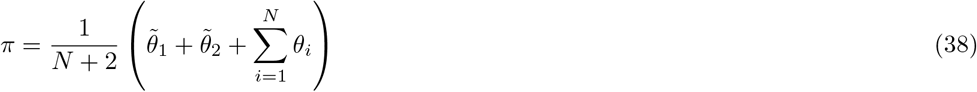

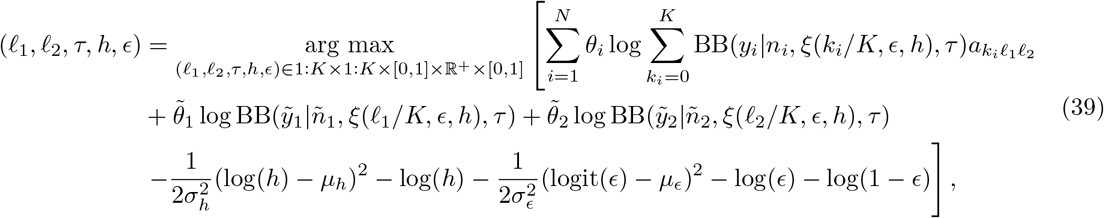

where

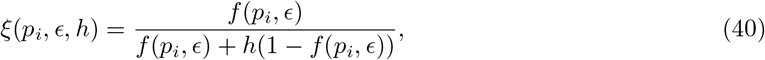

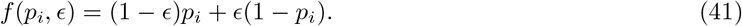

## B Theorems relating to preferential pairing

See Section 2.10 for notational definitions.

### Theorem 1.

*The number of configurations m given a K-ploid parent has ℓ copies of a are*

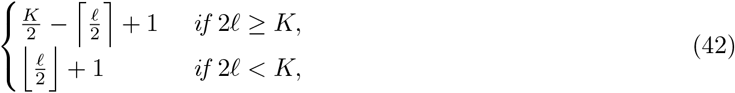

*where* ⌊·⌋ *and* ⌈·⌈ *are the ceiling and floor functions, respectively*.

### *Proof*.

We begin by placing as many a’s into an AA pair as possible. This is ⌊*ℓ*/2⌋ AA pairs. In this configuration, there are *K*/2 – ⌈*ℓ*/2⌉ aa pairs. We then navigate to a new configuration by subtracting 1 from *m*_1_ and *m*_3_ and adding 2 to *m*_2_. We do this until either *m*_1_ = 0 or *m*_3_ = 0. Proceeding in this way, we reach all possible configurations. Counting the number of configurations we obtained, we get (42).

Summing over *ℓ*, we can also get a result for the total number of possible values of m not conditional on *ℓ*.

### Theorem 2.

*The total number of possible values of **m** is*

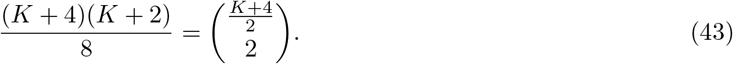

### *Proof*.

The proof involves summing (42) over *ℓ* under the two cases where *K*/2 is even and *K*/2 is odd.

We begin with the case where *K*/2 is even. Then the total number of elements is

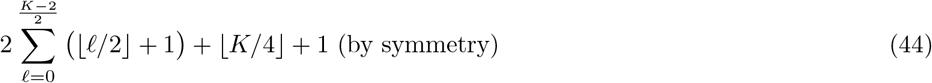

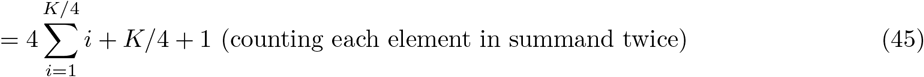

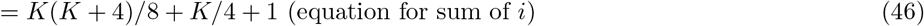

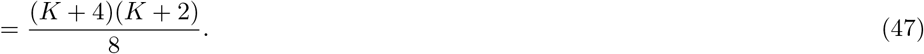

For the case where *K*/2 is odd, we have the total number of elements is

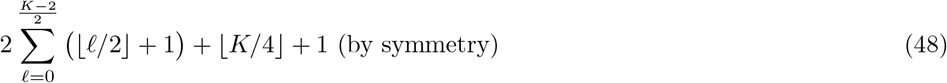

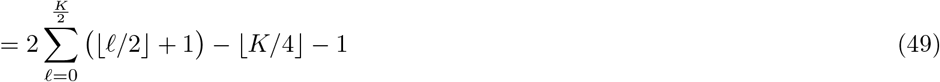

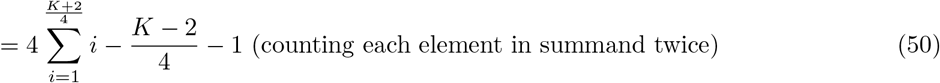

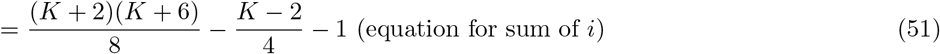

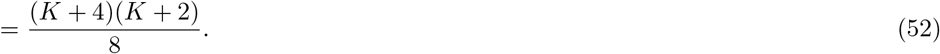

### Theorem 3.

*The total possible number of pairings where chromosomes are labeled but pairings are not labeled is*

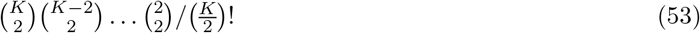

### *Proof*.

The proof is very similar the proof for the derivation of combinations. First, choosing 2 chromosomes from *K* is 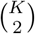, then choosing 2 chromosomes from *K* – 2 is 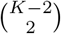. We continue in this manner until we exhaust all of the chromosomes. We then need to divide by the number of orderings for iteratively choosing 2 chromosomes, which is just 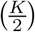!.

### Theorem 4.

*Let*

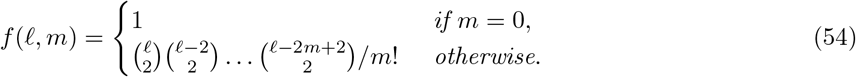

*Then the number of pairings that result in a configuration m where the chromosomes are labeled but the pairings are not labeled is*

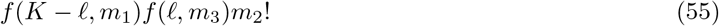

### *Proof*.

First, we select the number of pairings resulting in aa and we obtain *f*(*K* – *ℓ, m*_1_). The proof of this is very similar to the proof in Theorem 3 so we omit the details. Similarly, the number of pairings that result in AA is *f*(*ℓ, m*_3_). The number of ways we can pair the leftovers into Aa is *m*_2_! because we first choose one of *m*_2_ a’s to pair with the first a, then *m*_2_ – 1 of the remaining a’s to pair with the second a, and continue in this way until all a’s and a’s are paired. We then simply multiply these counts together to obtain (55).

### Theorem 5.

*If there is no preferential pairing, then the probability of configuration **m** is*

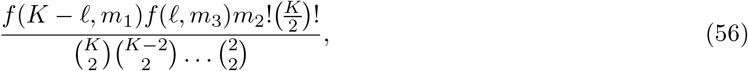

*where f*(*ℓ, m*) *is defined in* (54).

### *Proof*.

There are 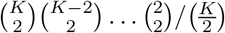! possible configurations where the chromosomes are labeled but the pairings are not labeled (Theorem 3). Of those, *f*(*K* – ℓ, m_1_)*f*(*ℓ,m*_3_)*m*_2_! contain configuration ***m*** (Theorem 4). If each configuration, where the chromosomes are labeled but the pairings are not, have an equal probability, then the probability of configuration ***m*** is just the ratio of these two quantities, (56).

## C Supplementary figures

**Figure S1:**
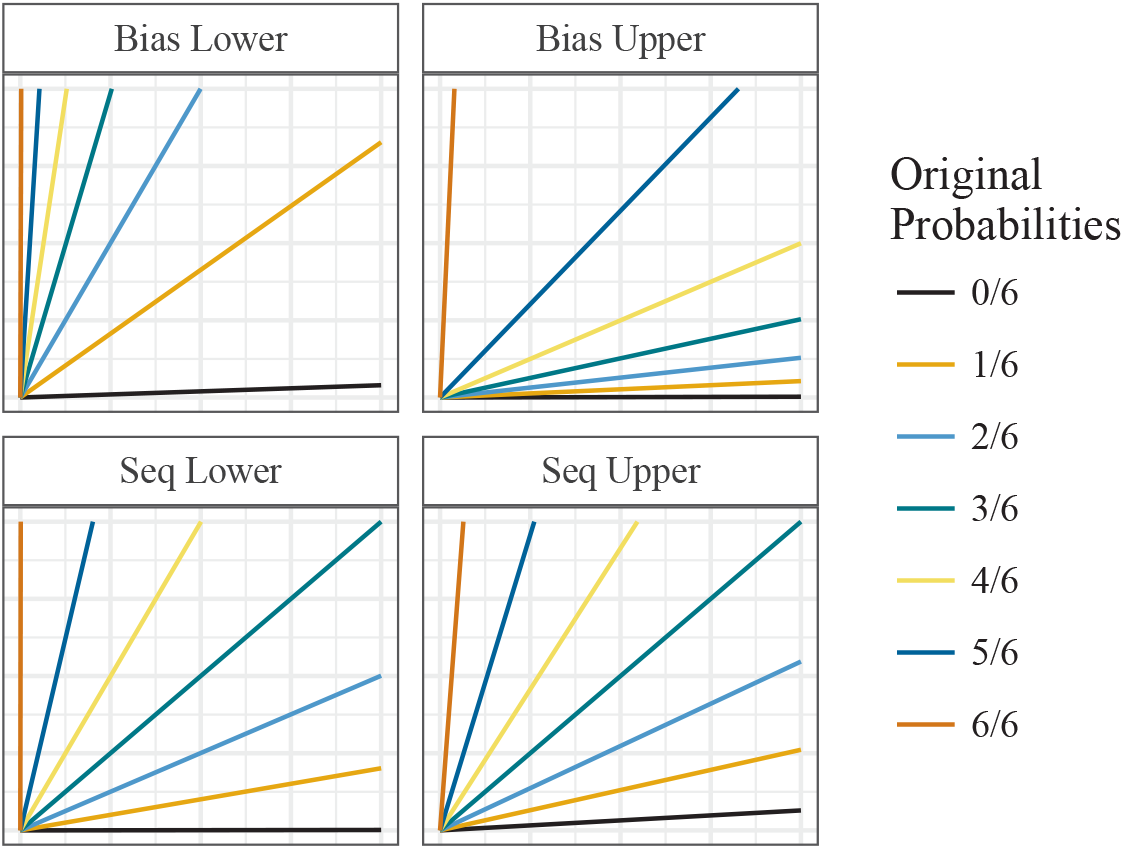
Considering an autohexaploid loci, −2 standard deviations of the bias parameter under our default prior (with a sequencing error rate of 0.01) (top left), +2 standard deviations of the bias parameter under our default prior (with a sequencing error rate of 0.01) (top right), −2 standard deviations of the sequencing error rate under our default prior (with a bias parameter of 1) (bottom left), +2 standard deviations of the sequencing error rate under our default prior (with a bias parameter of 1) (bottom right). The *x*-axis is the number of alternative reads and the *y*-axis is the number of reference reads.

**Figure S2:**
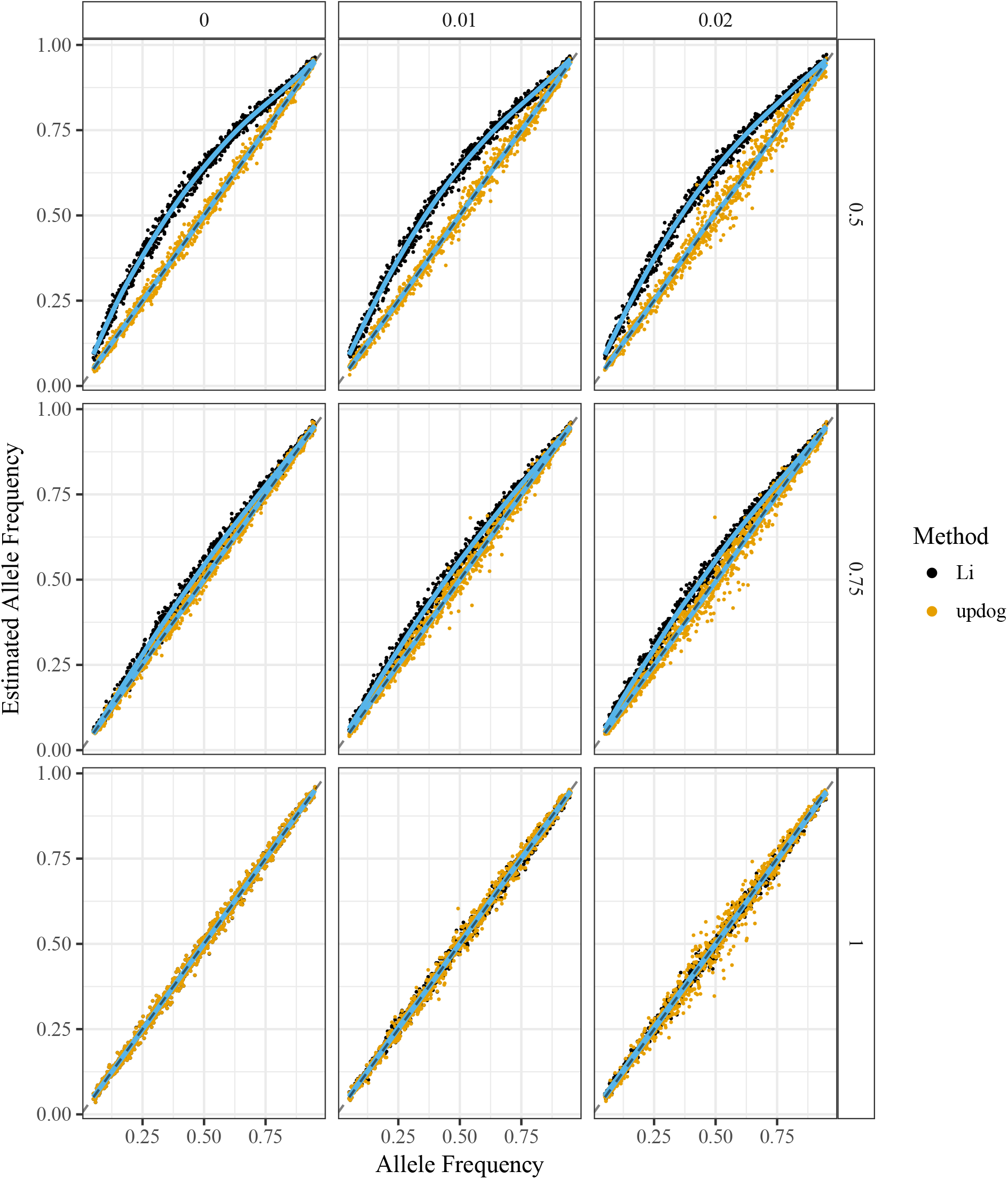
Estimated allele frequency (*y*-axis) for the `updog` method (black) and the method of Li [2011] (orange) versus true allele frequency (*x*-axis). An unbiased method would result in most points lying along the *y* = *x* line. A smooth generalized additive model was fit to the results of both methods (blue lines). The column facets distinguish between different levels of the overdispersion parameter and the row facets distinguish between different levels of the bias parameter (with 1 indicating no bias).

**Figure S2:**
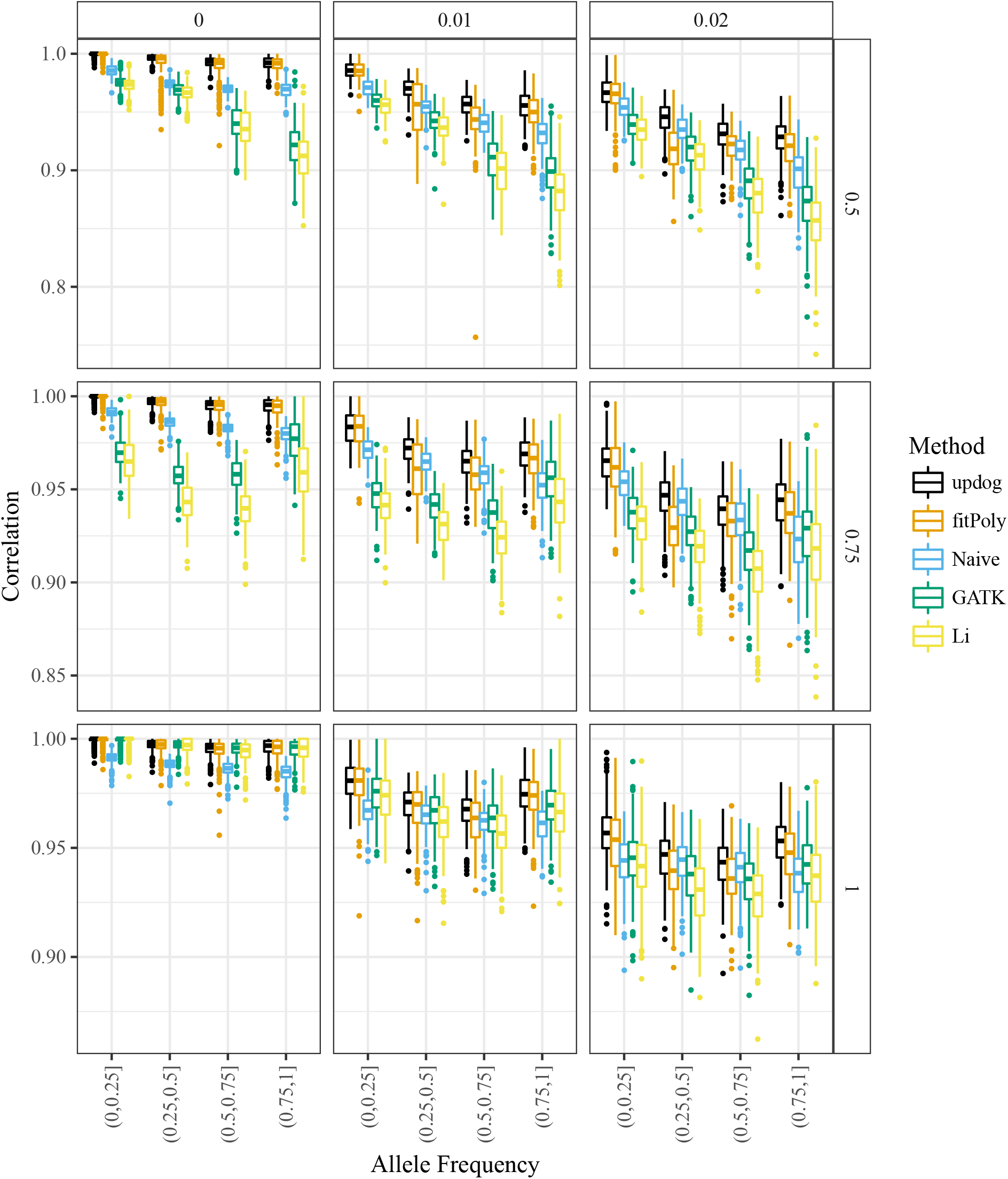
Boxplots of the correlation between the true and estimated genotypes (*y*-axis) stratified by the binned values of the major allele frequency (*x*-axis). The boxplots are color-coded by method: `updog` (black), `fitPoly` (orange), the naive method (blue), GATK (green), and the method of Li [2011] (yellow). The column facets distinguish between different levels of the overdispersion parameter and the row facets distinguish between different levels of the bias parameter (with 1 indicating no bias).

**Figure S3:**
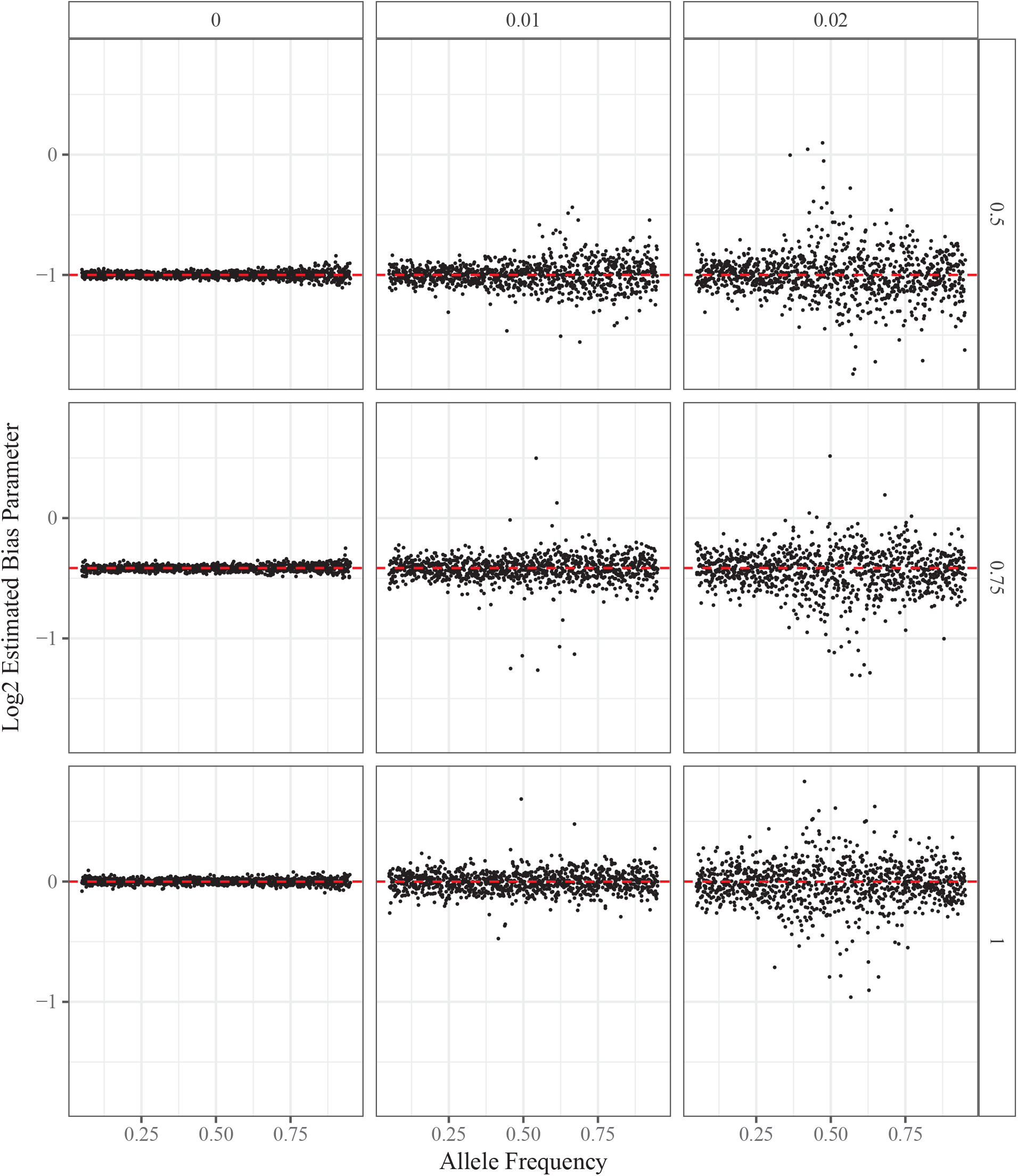
Allele frequency (*x*-axis) versus log_2_ of the estimated bias (*y*-axis). The row facets distinguish different levels of bias (with 1 indicating no bias) and the column facets distinguish different levels of overdispersion.

**Figure S4:**
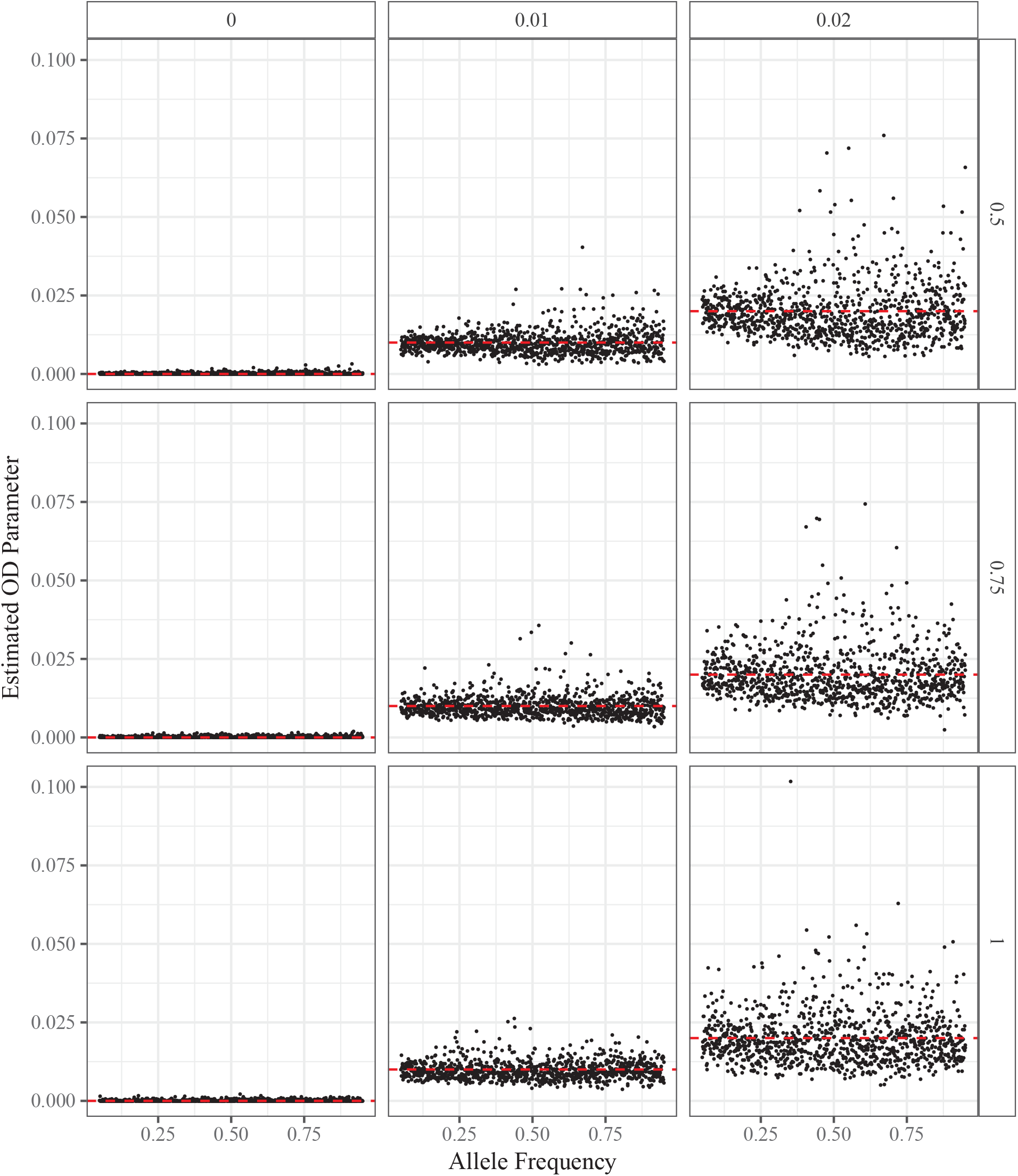
Allele frequency (*x*-axis) versus estimated overdispersion level (*y*-axis). The row facets distinguish different levels of bias (with 1 indicating no bias) and the column facets distinguish different levels of overdispersion.

**Figure S5:**
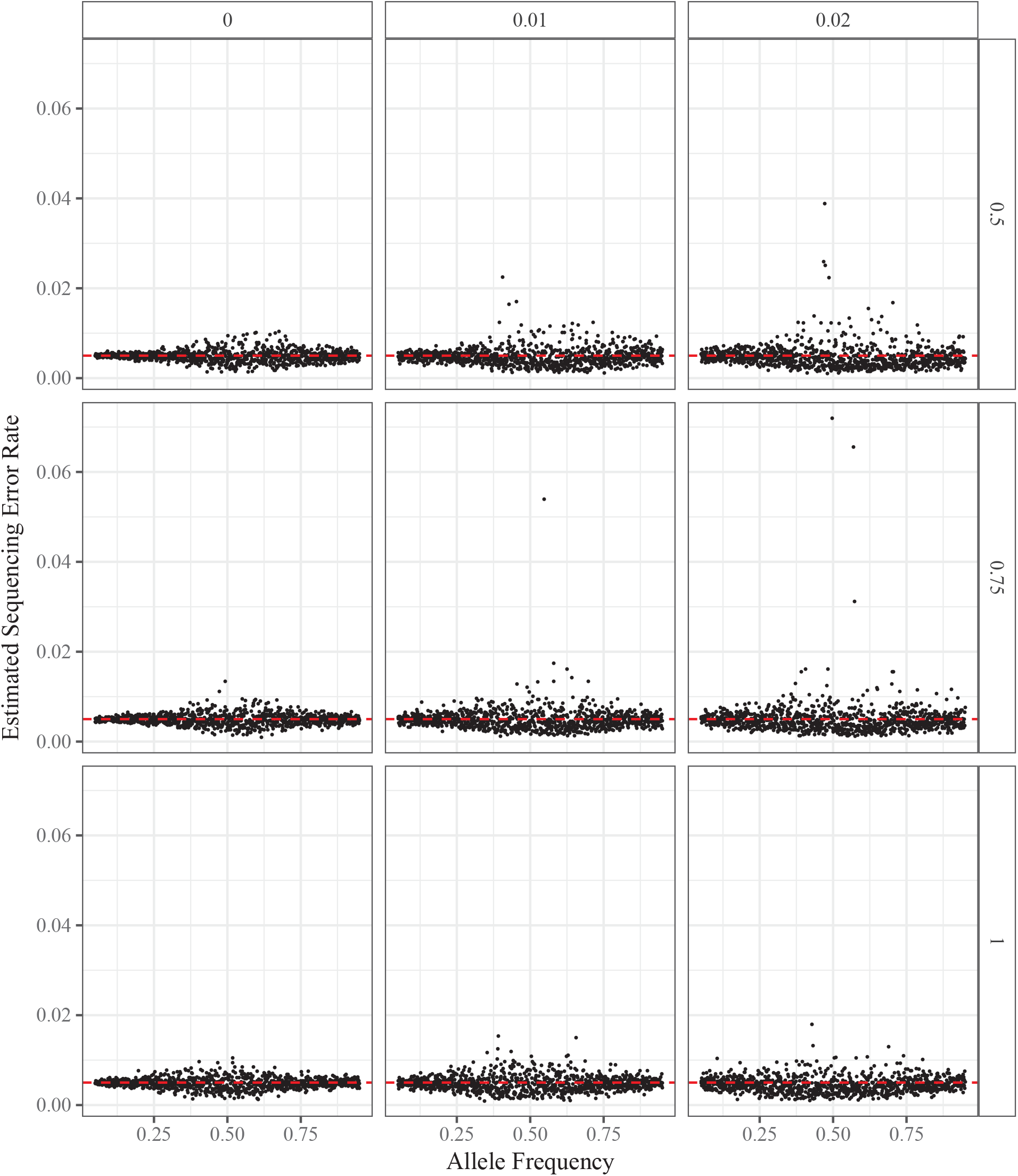
Allele frequency (*x*-axis) versus estimated sequencing error rate (*y*-axis). The row facets distinguish different levels of bias (with 1 indicating no bias) and the column facets distinguish different levels of overdispersion.

**Figure S6:**
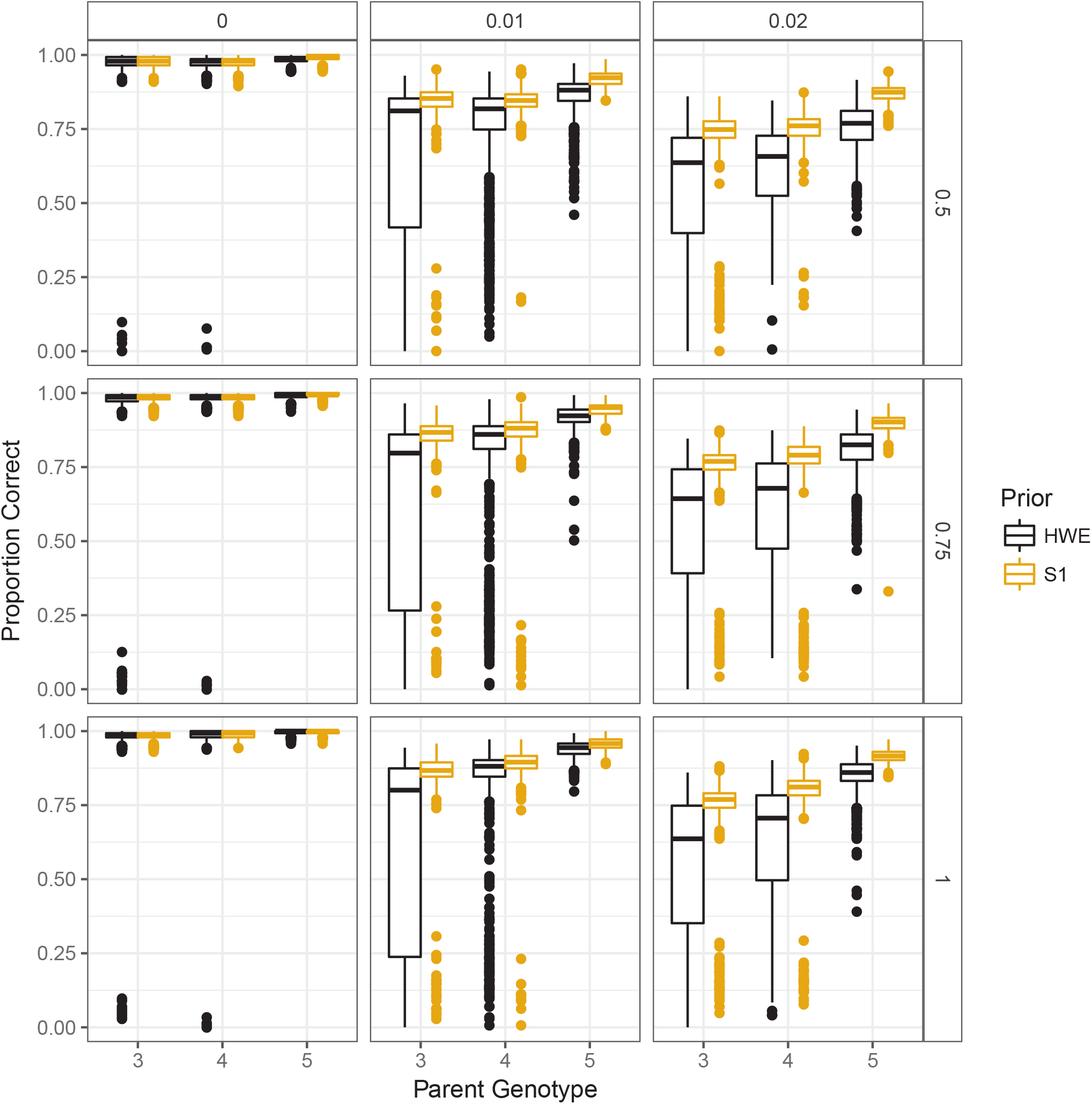
Box plots of the proportion of individuals genotyped correctly in an `updog` fit (*y*-axis) stratified by parent genotype (*x*-axis). Column facets distinguish different levels of overdispersion while row facets distinguish different levels of bias (with 1 indicating no bias). Orange box plots correctly assume the individuals are from an S1 population while black box plots incorrectly assume the population is in Hardy-Weinberg equilibrium.

**Figure S7:**
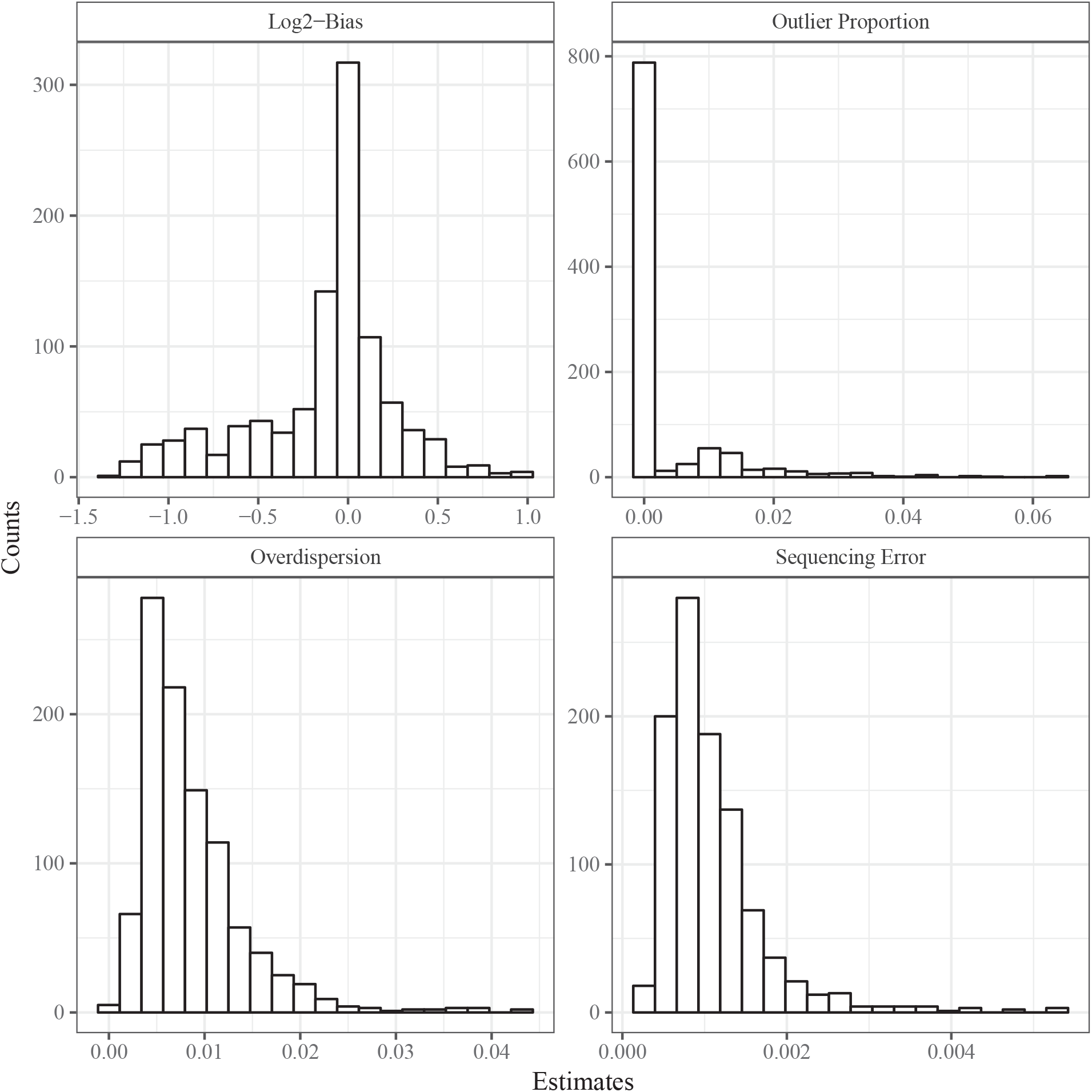
Histograms of `updog` estimates of parameters in 1000 SNPs from the data from Shirasawa et al. [2017].

**Figure S8:**
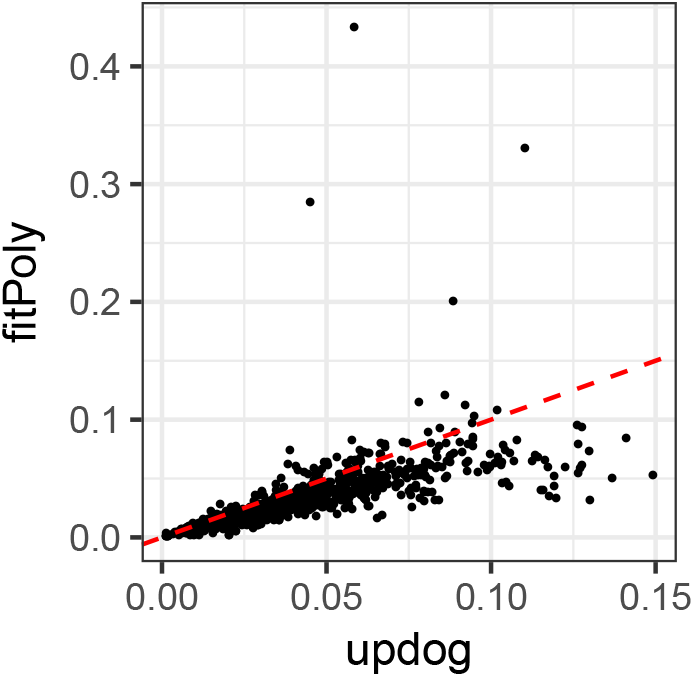
`Updog`’s estimated proportion of individuals genotyped incorrectly (*x*-axis) versus `fitPoly`’s estimated proportion of individuals genotyped incorrectly (*y*-axis) for the 1000 SNPs in the dataset from Shirasawa et al. [2017] with the largest read depth. The dashed horizontal line is the *y* = *x* line. `Updog` has a larger estimate of incorrectly genotyped individuals for points below this line, and a smaller estimate for points above this line.

**Figure S9:**
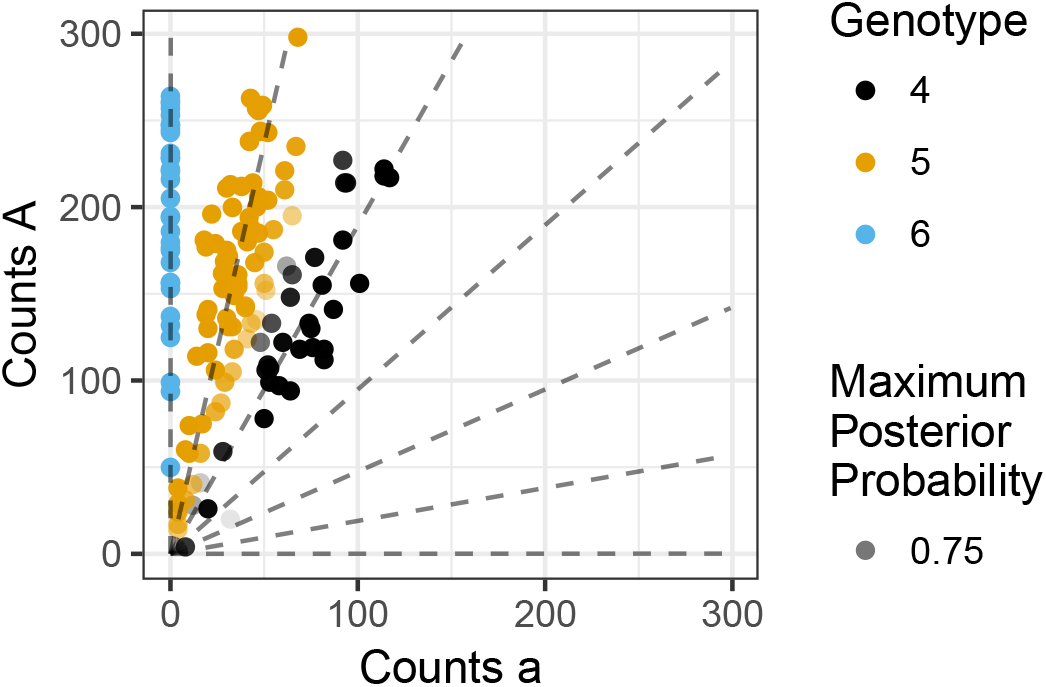
A genotype plot of the SNP from the dataset of Shirasawa et al. [2017] where `updog` estimated that 4.5% of the individuals were genotyped incorrectly and `fitPoly` estimated that 28.5% of the individuals were genotyped incorrectly. The figure is annotated by an `updog` fit.

**Figure S10:**
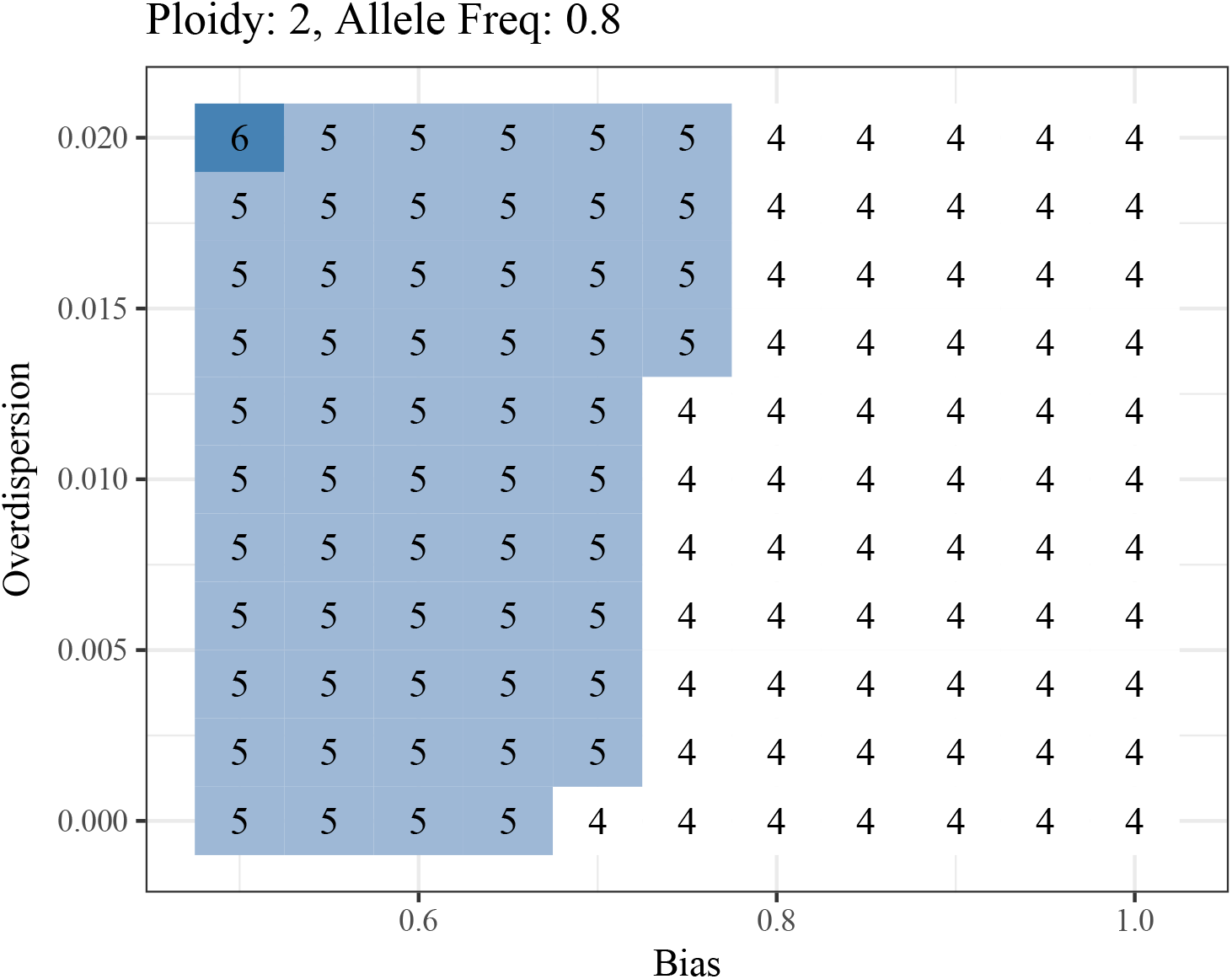
The minimum read depth required to obtain an oracle genotyping error rate less than 0.05 under different levels of overdispersion (*y*-axis) and allelic bias (*x*-axis) while the sequencing error rate is fixed at 0.001. This is for a diploid population under Hardy-Weinberg equilibrium with a major allele frequency of 0.8.

**Figure S11:**
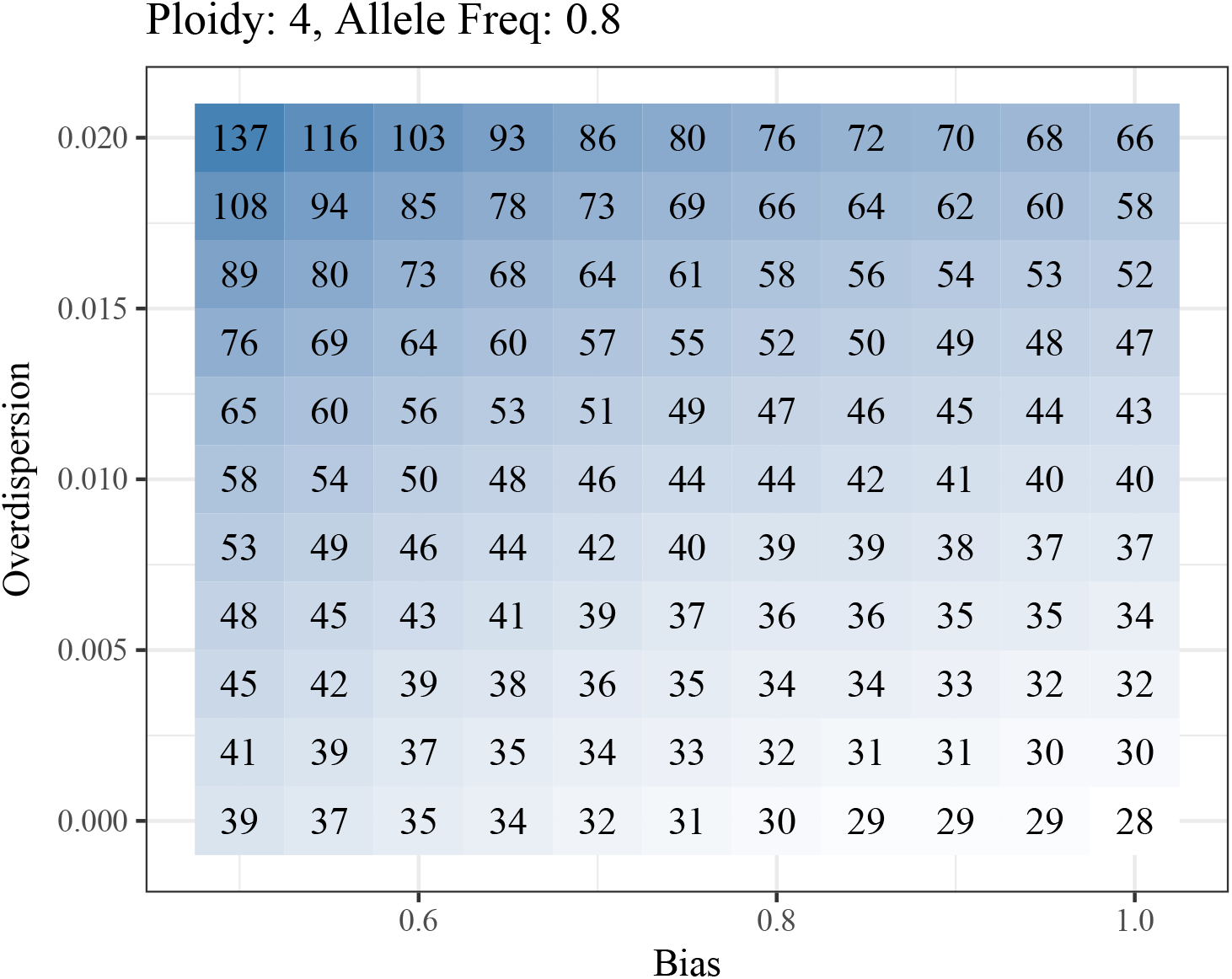
The minimum read depth required to obtain an oracle genotyping error rate less than 0.05 under different levels of overdispersion (*y*-axis) and allelic bias (*x*-axis) while the sequencing error rate is fixed at 0.001. This is for a tetraploid population under Hardy-Weinberg equilibrium with a major allele frequency of 0.8.

**Figure S12:**
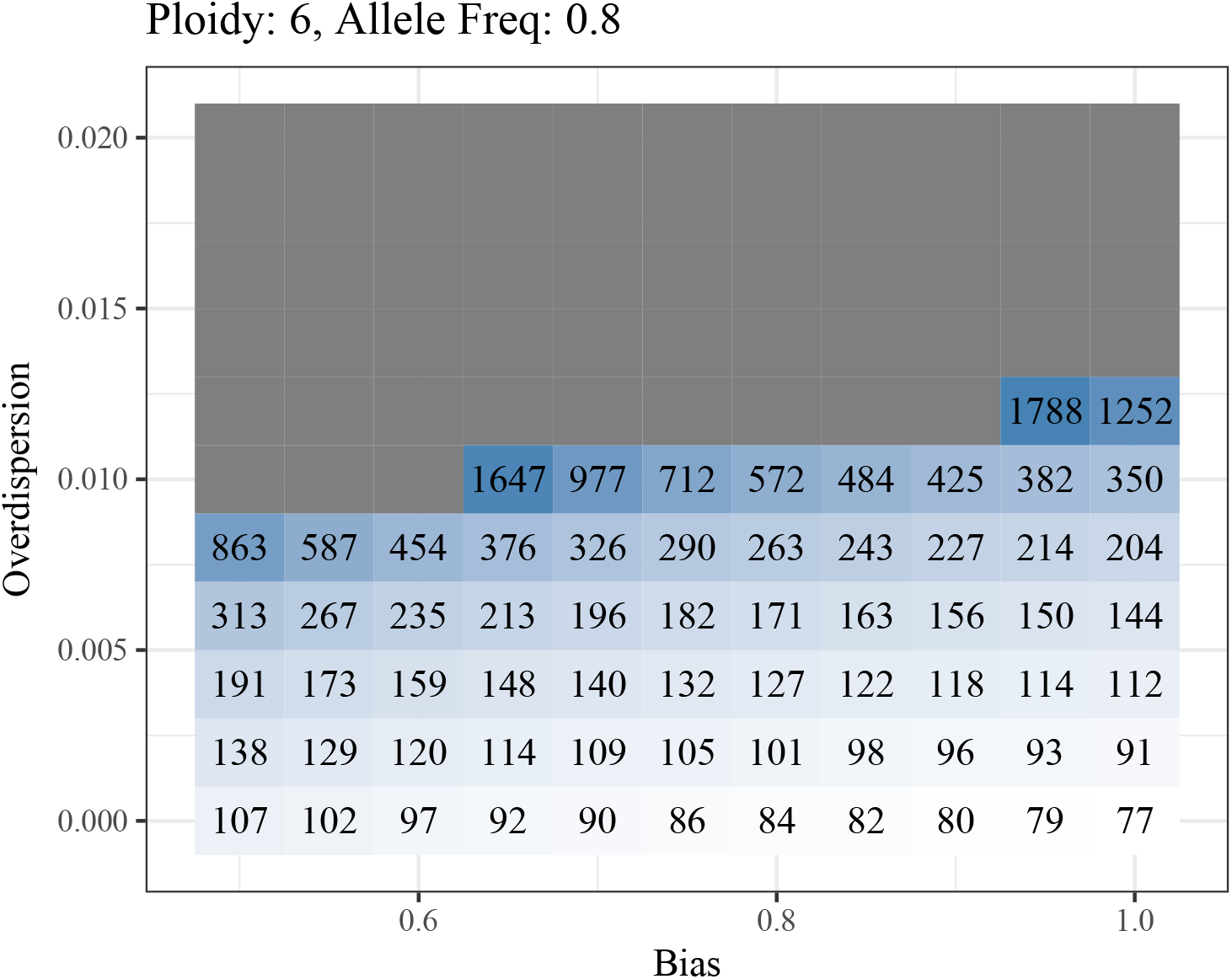
The minimum read depth required to obtain an oracle genotyping error rate less than 0.05 under different levels of overdispersion (*y*-axis) and allelic bias (*x*-axis) while the sequencing error rate is fixed at 0.001. This is for a hexaploid population under Hardy-Weinberg equilibrium with a major allele frequency of 0.8. Grey regions require a read depth greater than 3000.

**Figure S13:**
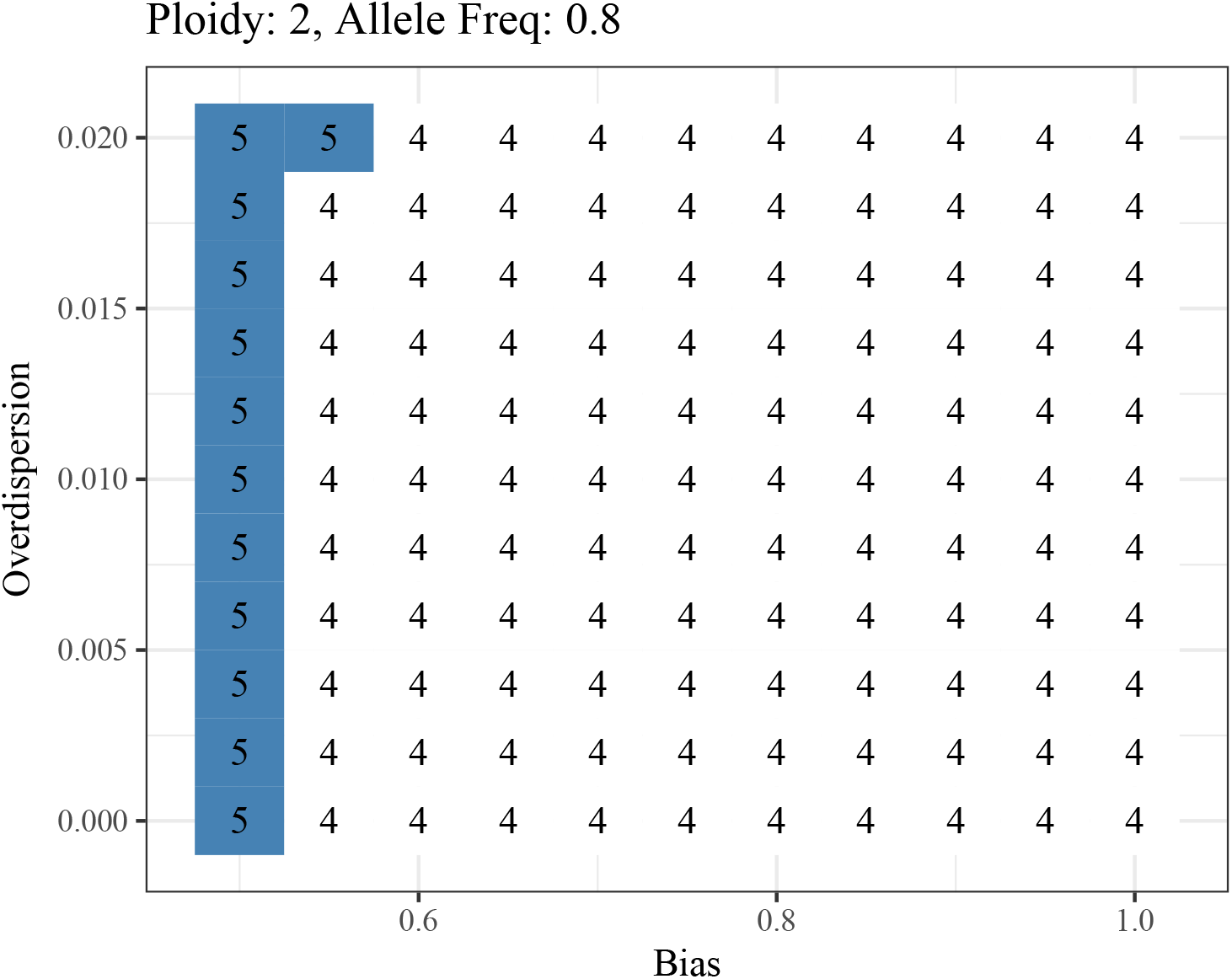
The minimum read depth required to obtain a correlation of over 0.9 between the true and oracly estimated genotypes under different levels of overdispersion (*y*-axis) and allelic bias (*x*-axis) while the sequencing error rate is fixed at 0.001. This is for a diploid population under Hardy-Weinberg equilibrium with a major allele frequency of 0.8.

**Figure S14:**
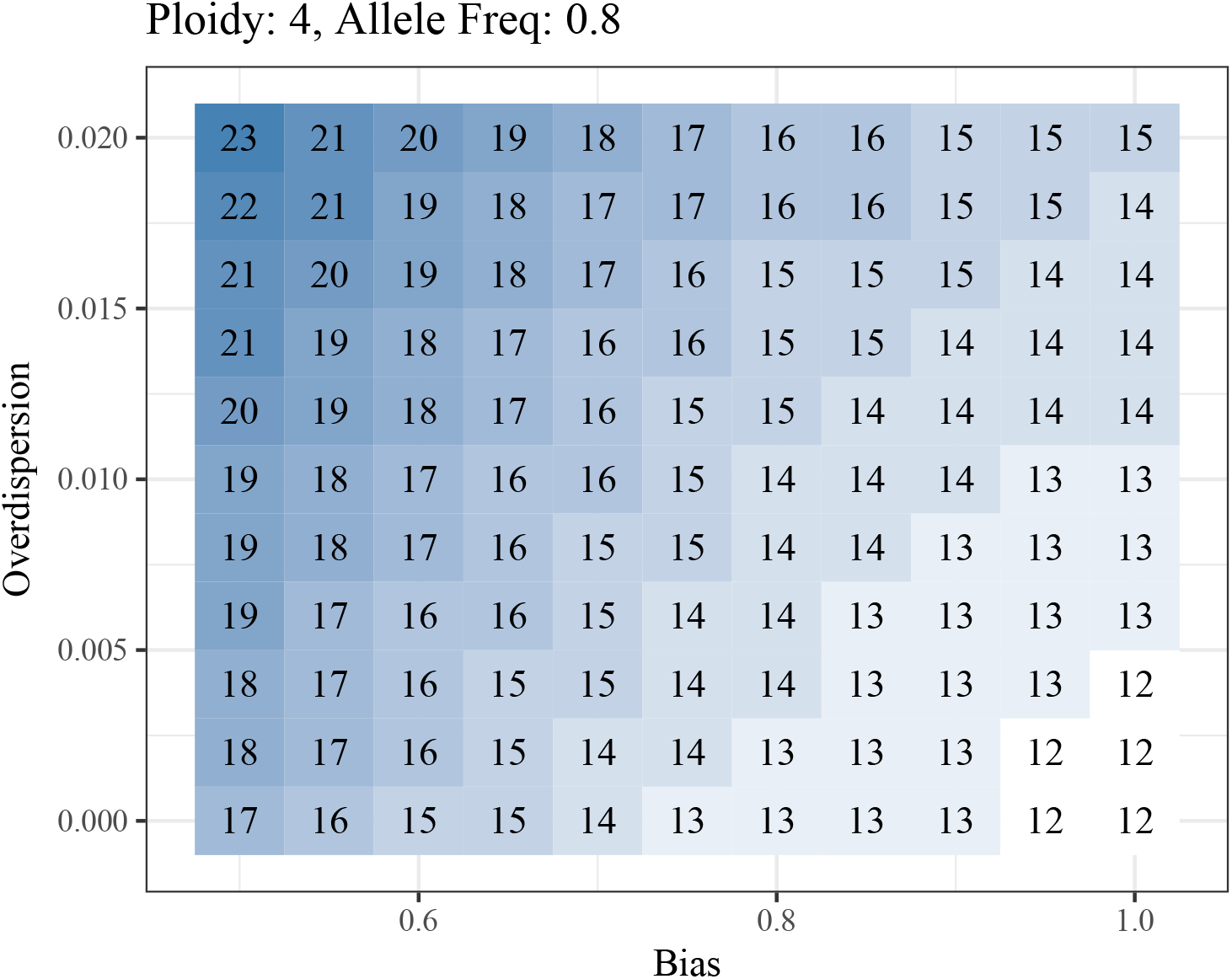
The minimum read depth required to obtain a correlation of over 0.9 between the true and oracly estimated genotypes under different levels of overdispersion (*y*-axis) and allelic bias (*x*-axis) while the sequencing error rate is fixed at 0.001. This is for a tetraploid population under Hardy-Weinberg equilibrium with a major allele frequency of 0.8.

**Figure S15:**
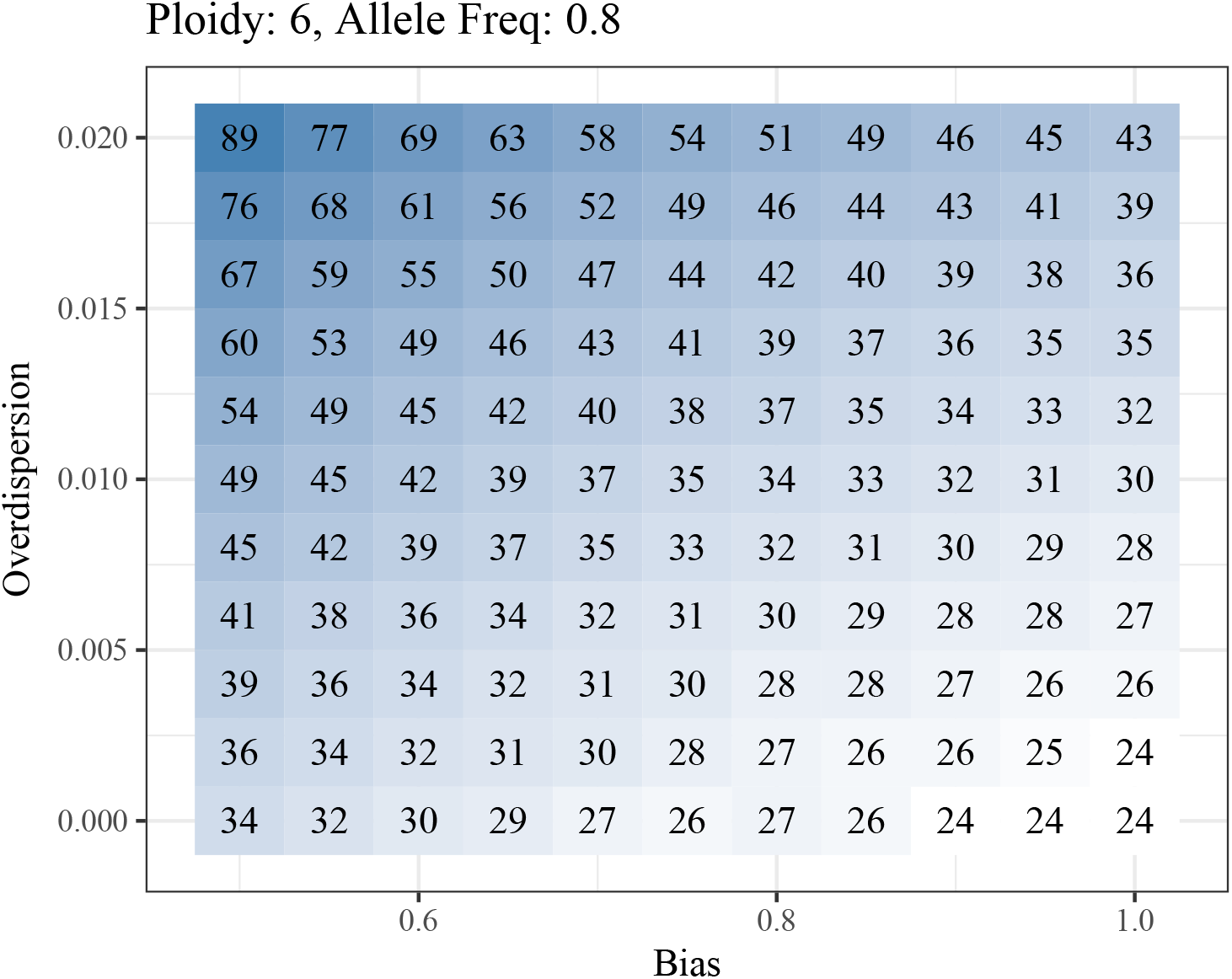
The minimum read depth required to obtain a correlation of over 0.9 between the true and oracly estimated genotypes under different levels of overdispersion (*y*-axis) and allelic bias (*x*-axis) while the sequencing error rate is fixed at 0.001. This is for a hexaploid population under Hardy-Weinberg equilibrium with a major allele frequency of 0.8.

